# A suppressor of axillary meristem maturation promotes longevity in flowering plants

**DOI:** 10.1101/2020.01.03.893875

**Authors:** Omid Karami, Arezoo Rahimi, Majid Khan, Marian Bemer, Rashmi R. Hazarika, Patrick Mak, Monique Compier, Vera van Noort, Remko Offringa

## Abstract

Post embryonic development and growth of flowering plants are for a large part determined by the activity and maturation state of stem cell niches formed in the axils of leaves, the so-called axillary meristems (AMs)^1,2^. Here we identify a new role for the Arabidopsis *AT-HOOK MOTIF CONTAINING NUCLEAR LOCALIZED 15* (*AHL15*) gene as a suppressor of AM maturation. Loss of *AHL15* function accelerates AM maturation, whereas ectopic expression of *AHL15* suppresses AM maturation and promotes longevity in Arabidopsis and tobacco. Together our results indicate that *AHL15* expression acts as a key molecular switch, directly downstream of flowering genes (*SOC1, FUL*) and upstream of GA biosynthesis, in extending the plant’s lifespan by suppressing AM maturation.

---

Plant architecture and -longevity are dependent on the activity of stem cell groups called meristems. The primary shoot and root apical meristem of a plant are established during early embryogenesis and give rise to respectively the shoot- and the root system during post-embryonic development. In flowering plants, post-embryonic shoot development starts with a vegetative phase, during which the primary shoot apical meristem (SAM) produces morphogenetic units called phytomers consisting of a stem (internode) subtending a node with a leaf and a secondary or axillary meristem (AM) located in the axil of the leaf^1,2^. Both the SAM and these AMs undergo a maturation process. Like the SAM, young immature AMs are vegetative and when activated they produce leaves, whereas partially matured AMs produce a few cauline leaves before they fully mature into inflorescence meristems (IMs) and start developing phytomers comprising a stem subtending one or more flowers^2,3^.

The rate of AM maturation is an important determinant of plant longevity and -life history. Monocarpic plants, such as the annuals *Arabidopsis thaliana* (Arabidopsis) or *Nicotiana tabacum* (tobacco), complete their life cycle in a single growing season. The immature AMs that are established during the vegetative phase initially produce leaves. Upon floral transition, however, all AMs rapidly mature into IMs, producing secondary and tertiary inflorescences with bracts and flowers, thus maximizing offspring production before the plant’s life ends with senescence an death. The number of leaves and bracts produced by an AM is thus a measure for its maturation state upon activation. By contrast, many other flowering plant species are polycarpic, such as the close Arabidopsis relative *Arabidopsis lyrata*. Under permissive growth conditions, they can live and flower for more than two growing seasons. As some AMs are maintained in the immature vegetative state, this allows polycarpic plants to produce new shoots after seed set or during the next growing season^4,5^. Despite considerable interest in the molecular basis of plant life history, the proposed molecular mechanisms determining the difference in AM maturation between monocarpic and polycarpic plants are still largely based on our extensive knowledge on the control of flowering in Arabidopsis and closely related species. From these studies, the MADS box proteins SUPPRESSOR OF OVEREXPRESSION OF CONSTANS 1 (SOC1) and FRUITFULL (FUL) have been identified as key promotors of flowering and monocarpic growth, and FLOWERING LOCUS C (FLC) as their upstream cold-sensitive inhibitor^5–7^. However, the factors that suppress AM maturation, and thereby promote polycarpic growth, are still elusive.

Here we present evidence that the Arabidopsis *AT-HOOK MOTIF COINTAINING NUCLEAR LOCALIZED 15* (*AHL15*) gene is a key switch in the control of AM maturation. The gene was initially selected from a yeast one-hybrid screen, and was shown to form a clade with 14 other *AHL* genes in Arabidopsis that encode nuclear proteins containing a single N-terminal DNA binding AT-hook motif and a C-terminal Plants and Prokaryotes Conserved (PPC) domain (**Supplementary Fig. 1a**). The PPC domain was previously shown to contribute to the physical interaction with other AHL or nuclear proteins^8^. *AHL15* homologs have been implicated in several aspects of plant growth and development in Arabidopsis, including hypocotyl growth and leaf senescence ^8–10^, flower development^11^, and flowering time^10,12^.

In contrast to other *ahl* mutants^10^, *ahl15* loss-of-function mutant plants (**Supplementary Fig. 1b-e**) flowered at the same time and developed the same number of rosette leaves before flowering as wild-type plants under short day (SD) and log day (LD) conditions (**Supplementary Fig. 2d,e**). After bolting, however, the AMs located in the axils of rosette leaves (rosette AMs) of *ahl15* mutant plants produced less additional rosette leaves compared to those in wild-type plants (**Fig. 1a,e and Supplementary Fig. 3a-c)**. More detailed analysis showed that this reduction in additional rosette leaf production in *ahl15* plants was not caused by a delayed outgrowth of rosette AMs into axillary buds or to early floral transition, but rather to a reduction in the vegetative activity of these buds (**Supplementary Fig. 4a,b**). Following the floral transition, the rosette AMs did produce the same number cauline leaves (**Supplementary Fig. 3d**) and flowers (**Supplementary Fig. 5a**) in wild type (Col-0) and the *ahl15* mutant. However, the cauline branches produced by the aerial AMs on *ahl15* inflorescences developed less cauline leaves (**Supplementary Fig. 3e**) and flowers/fruits (**Supplementary Fig. 5b**) compared to those on wild-type inflorescences, resulting in a significant reduction of the total number of cauline leaves and flowers on *ahl15* inflorescences. Introduction of the *pAHL15:AHL15* genomic clone into the *ahl15* mutant background restored both the rosette and cauline leaf and flower and fruit numbers to wild-type levels (**Fig. 1e and Supplementary Fig. 3a-e, Supplementary Fig. 5b**), confirming that the phenotypes were caused by *ahl15* loss-of-function. GUS staining of plants carrying a *pAHL15:GUS* promoter:reporter fusion showed that *AHL15* is expressed in AMs (**Fig. 1a,b**) and young axillary buds (**Fig. 1d**). Together these results suggested a novel role for *AHL15* in controlling AM maturation.

**Fig. 1.**
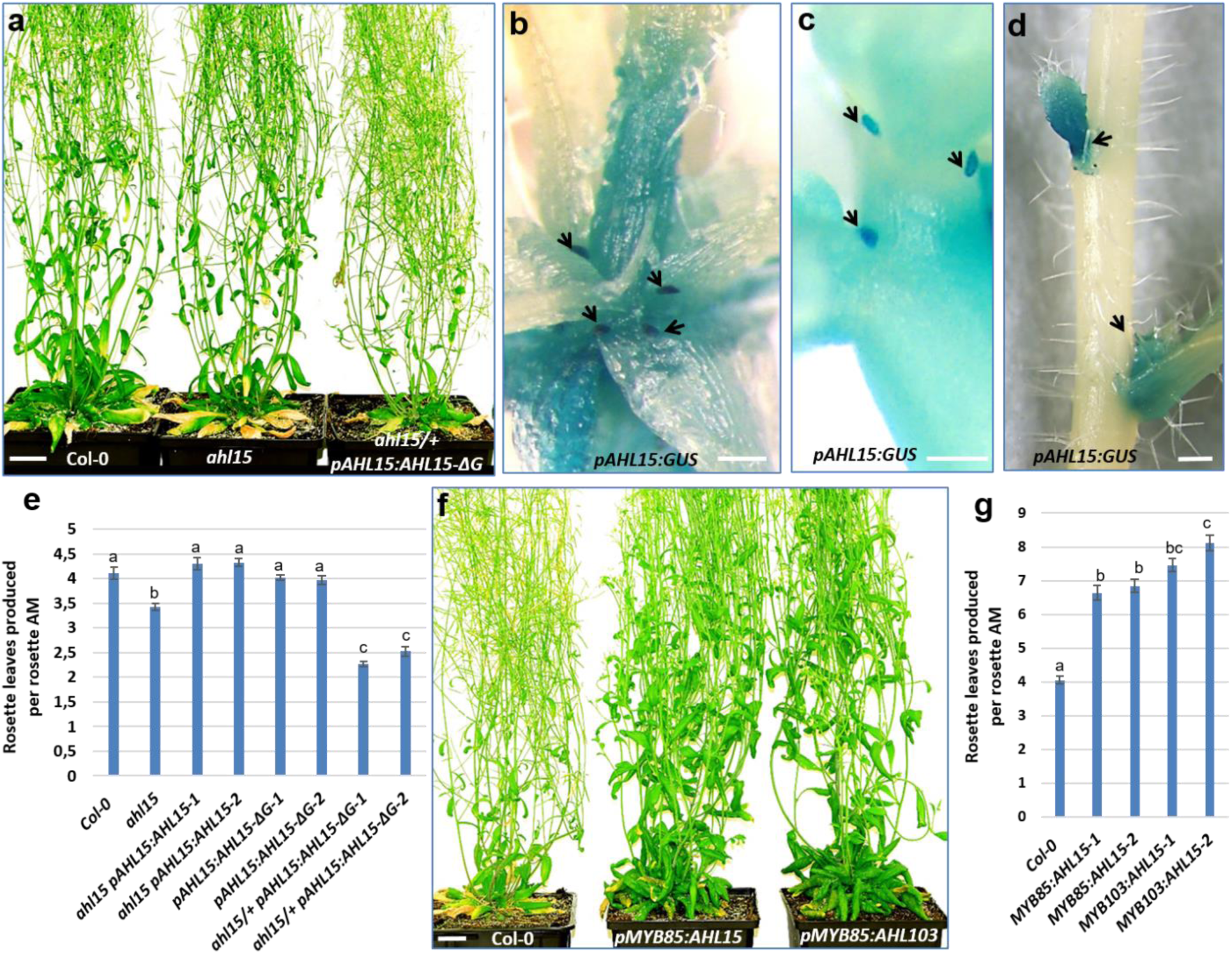
*AHL15* represses AM maturation in Arabidopsis. (**a**) Shoot phenotypes of fifty-day-old flowering wild-type (left), *ahl15* (middle) and *ahl15/+ pAHL15:AHL15-ΔG* mutant (right) plants. (**b**-**d**) *pAHL15:GUS* expression in rosette AMs (arrow heads in **b**), aerial AMs located on an young inflorescence stem (arrow heads in **c**) and in activated axillary buds on an inflorescence stem (arrow heads in **d**) of a flowering plant. (**e**) The rosette leavesproduced per rosette AM in fifty-day-old wild-type, *ahl15, ahl15 pAHL15:AHL15, pAHL15:AHL15-ΔG* and *ahl15/+ pAHL15:AHL15-ΔG* plants (n=15). (**f**) Shoot phenotype of a sixty-day-old flowering wild-type (left), *pMYB85:AHL15* (middle) or *pMYB103:AHL15* (right) plant. (**g**) The rosette leaves produced per rosette AM in sixty-day-old wild-type, *pMYB85:AHL15* or *pMYB103:AHL15* plants (n=15). Error bars in **e** and **g** represent the standard error of the mean. Letters (a, b, c) indicate statistically significant differences, as determined by a one-way ANOVA with a Tukey’s HSD post hoc test (p < 0.01). Plants were grown under long day conditions (LD). Size bars indicate 2 cm in **a, f** and **1** mm in **b-c**. The leaf production per rosette AM in **e** and **g** was determined for two independent transgenic lines (1 and 2).

AHL proteins interact with each other through their PPC domain and with other non-AHL proteins through a conserved six-amino-acid (GRFEIL) region in the PPC domain. Expression of an AHL protein without the GRFEIL region leads to a dominant negative effect, as it generates a non-functional complex that is unable to modulate transcription ^8^. Expression of a deletion version of AHL15 lacking the GRFEIL region under control of the *AHL15* promoter (*pAHL15:AHL15-ΔG*) in the wild-type background (n=20) resulted in fertile plants (**Supplementary Fig. 2a,b**) that showed normal AM maturation (**Fig. 1e and Supplementary Fig. 3**). In the heterozygous *ahl15* loss-of-function background, however, *pAHL15:AHL15-ΔG* expression induced early flowering (**Supplementary Fig. 2d,e and Supplementary Fig. 4b**), resulting in a strong reduction of rosette and cauline leaf production by AMs (**Fig. 1a,e, Supplementary Fig. 3a-f, Supplementary Fig. 4a**). Homozygous *ahl15 pAHL15:AHL15-ΔG* progeny were never obtained, and defective seeds present in siliques of *ahl15/+ pAHL15:AHL15-ΔG* plants suggest that this genetic combination is embryo lethal (**Supplementary Fig. 2c**). The significantly stronger phenotypes observed for *ahl15/+ pAHL15:AHL15-ΔG* plants are in line with the dominant negative effect of AHL15-ΔG expression overcoming the functional redundancy among Arabidopsis clade A *AHL* family members ^8–10^.

Based on the observation that the flowering time and the number of rosette leaves before bolting was the same for wild-type and *ahl15* loss-of-function plants, but not in *ahl15/+ pAHL15:AHL15-ΔG* plants, we speculated that other *AHL* clade A family members are more active in the SAM, whereas *AHL15* more strongly acts on AM maturation. To test this, we overexpressed a fusion protein between AHL15 and the rat glucocorticoid receptor under control of the constitutive Cauliflower Mosaic Virus (CaMV) *35S* promoter (*p35S:AHL15-GR*). This rendered the nuclear import and thus the activity of the ectopically expressed AHL15-GR fusion inducible by dexamethasone (DEX). Untreated *p35S:AHL15-GR* plants showed a wild-type phenotype (**Supplementary Fig. 6b,c**), but after spraying flowering *p35S:AHL15-GR* plants with DEX, rosette AMs produced significantly more rosette and cauline leaves (**Supplementary Fig. 6a-c**). Interestingly, spraying *p35S:AHL15-GR* plants before flowering also significantly delayed their floral transition (**Supplementary Fig. 6d**), indicating that ectopically expressed AHL15 can also suppress maturation of the SAM. In turn, overexpression of the Arabidopsis *AHL* family members *AHL19, AHL20, AHL27* and *AHL29*, as well as the putative *AHL15* orthologs from *Brassica oleracea* and *Medicago truncatula* in Arabidopsis resulted in similar morphological changes as observed for *p35S:AHL15-GR* plants after DEX treatment. The overexpression plants produced more rosette and cauline leaves during flowering (**Supplementary Fig. 8**), supporting the idea that there is functional redundancy among AHL clade A family members and that the ability to control either SAM or AM maturation depends on the spatio-temporal expression of the corresponding genes.

In contrast to the observed growth arrest and death of two-month-old Arabidopsis plants grown under long day (LD) conditions (**Fig. 2d**), four-month-old short day (SD) grown Arabidopsis plants continued to grow after the first cycle of flowering, as aerial AMs on the last-formed lateral branches produced new rosette leaves (**Fig. 2a and Supplementary Fig. 9)**. However, SD grown *ahl15* mutant plants did not show this renewed vegetative growth and died, whereas *ahl15 pAHL15:AHL15* plants grew like wild type under these conditions (**Fig. 2a and Supplementary Fig. 9**). GUS staining of *pAHL15:GUS* plants revealed that *AHL15* expression was strongly enhanced in young lateral inflorescences, axils of cauline leaves and rosette branches in SD conditions compared to LD conditions (**Fig. 2b and Supplementary Fig. 10**), indicating that *AHL15* expression is day-length sensitive, and confirming a key role for this gene in suppressing AM maturation under SD conditions.

**Fig. 2.**
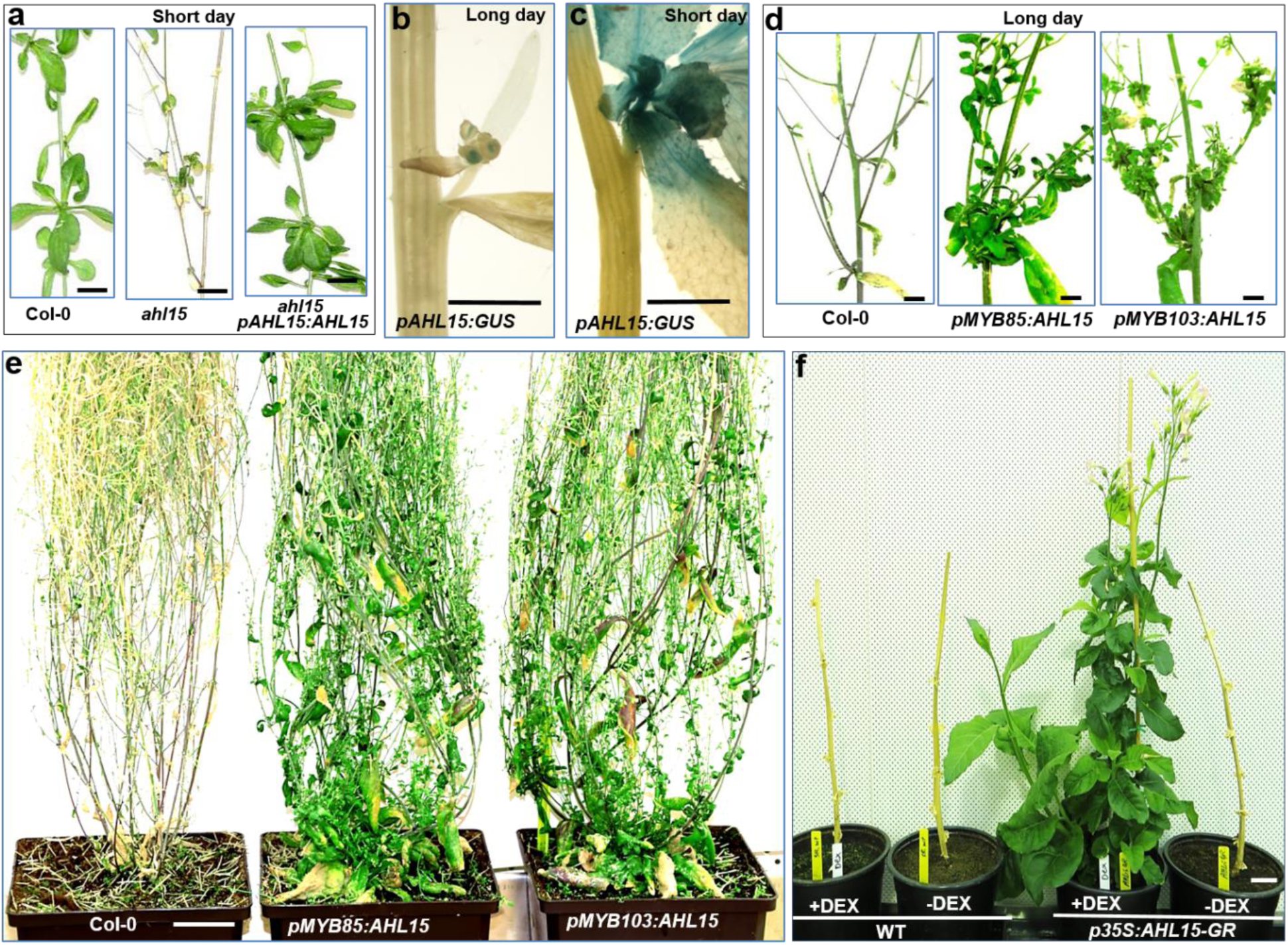
*AHL15* promotes longevity in Arabidopsis and tobacco. (**a**) Rosette leaves produced by aerial AMs in four-month-old wild-type (left) or *ahl15 pAHL15:AHL15* (right) plants, but not in *ahl15* mutant plants (middle), grown in short day (SD) conditions. (**b, c**) *pAHL15:GUS* expression in a lateral inflorescence of a 9-week-old plant grown in long day (LD) conditions (**b**), or a four-month-old plant grown in SD conditions (**c**). (**d**) Lateral aerial nodes without and with rosette leaves in four-month-old wild-type (left), *pMYB85:AHL15* (middle) or *pMYB103:AHL15* (right) plants grown in LD conditions. (**e**) Phenotype of a four-month-old wild-type (Col-0, left), *pMYB85:AHL15* (middle) or *pMYB103:AHL15* (right) plant grown in LD conditions. (**f**) Growth response following DEX treatment of bare stems of five-month-old wild-type (WT, left) or *35S:AHL15-GR* (right) tobacco plants grown in LD conditions. Size bars indicate 1 cm in **a-d** and 2 cm in **b** and 5 cm in **c**.

Growing Arabidopsis plants under SD conditions significantly delays flowering ^13^, and this might thus indirectly enhance the repressing effect of AHL15 on AM maturation. In order assess AHL15 function independently of day length and flowering time, we expressed *AHL15* under control of the *MYB85* or *MYB103* promoters, which are highly active in Arabidopsis rosette nodes and aerial axillary buds (**Supplementary Fig. 7a**) ^14^. In LD conditions, *pMYB103:AHL15* and *pMYB85:AHL15* plants flowered at the same time as wild-type plants, but their AMs produced significantly more rosette and cauline leaves compared to those in wild-type plants (**Fig. 1f,g and Supplementary Fig. 7b,c**). Moreover, after flowering and seed set, when wild-type plants senesced and died, *pMYB103:AHL15* and *pMYB85:AHL15* rosette and aerial AMs produced new rosette leaves, which allowed these plants to continue to grow and generate new flowers and seeds (**Fig. 2d,e**). Also senesced *p35S:AHL15-GR* plants carrying fully ripened siliques started new aerial vegetative development on lateral secondary inflorescences after DEX treatment, and ultimately produced new inflorescences from the resulting rosettes (**Supplementary Fig. 11a**). Interestingly, the development of vegetative shoots from AMs formed on rosette and aerial nodes after reproduction also contributes to the polycarpic growth habit of *Arabis alpina* plants^19^. Our results indicate that increased expression of *AHL15* in late stages of development promotes longevity by inducing a polycarpic-like growth habit in Arabidopsis.

To determine whether heterologous *AHL15* expression could induce similar developmental changes in a monocarpic plant species from a different family, we introduced the *35S:AHL15-GR* construct into tobacco. Wild-type and *p35S:AHL15-GR* tobacco plants were allowed to grow and set seeds without DEX treatment. After seed harvesting, all leaves and side branches were removed, and the bare lower parts of the primary stems were either mock- or DEX treated. Whereas stems of wild-type and mock-treated *p35S:AHL15-GR* plants did not show any growth, the AMs on DEX-treated *p35S:AHL15-GR* tobacco stems resumed vegetative growth, eventually leading to a second cycle of flowering and seed set (**Fig. 2e**). Continued DEX treatment after each subsequent cycle of seed harvesting efficiently induced vegetative growth and subsequent flowering and seed set, allowing the *p35S:AHL15-GR* tobacco plants to survive for more than 3 years (**Supplementary Fig. 11b**). This result confirmed the conclusion from previous overexpression experiments (**Supplementary Fig. 8**) that enhanced *AHL15* expression facilitates polycarpic-like growth by delaying AM maturation.

Previously, an Arabidopsis double loss-of-function mutant of the *SOC1* and *FUL* genes was found to show polycarpic-like growth as a result of reduced AM maturation ^6^. We found that the aerial rosette formation that is normally observed in the *soc1 ful* double mutant was significantly reduced in *soc1 ful ahl15* triple mutant plants (**Fig. 3a, b**). Moreover, the polycarpic-like features of the *soc1 ful* double mutant were lost in the *ahl15* mutant background, as *soc1 ful ahl15* plants senesced and died following seed set just like wild-type Arabidopsis. Expression analysis by qRT-PCR (**Fig. 3c**) or by using the *pAHL15:GUS* reporter (**Fig. 3d,e**) showed that *AHL15* was indeed strongly upregulated in *soc1 ful* inflorescence nodes and lateral inflorescences. Previous studies have revealed that the expression of *SOC1* is positively regulated by LD conditions. Moreover, SOC1 was shown to bind to the *AHL15* upstream and downstream regions (Chr3, 20603158-20604316 and 20610947-20612012), which both contain a canonical CArG box (CC[A/T]6GG] (**Fig. 3f**)^17–19^. This together with our own data suggested that *AHL15* expression is repressed by SOC1 in LD conditions (**Fig. 2c**), and that unrepressed AHL15 activity in the *soc1 ful* background explains the aerial rosette formation and the polycarpic-like growth of mutant plants. To check whether the CArG-box containing regions could also be bound by FUL, we used stem fragments containing axillary nodes of *pFUL:FUL-GFP ful* plants to perform ChIP. Subsequent qPCR revealed significant enrichment for the upstream and downstream regions (**Fig. 3g**), indicating that FUL can repress *AHL15* expression by directly binding to these regions. To further confirm that FUL and SOC1 can bind the CArG-boxes in the *AHL15* up- and downstream regions, we performed an Electrophoretic Mobility Shift Assay (EMSA) experiment. Probe fragments containing the corresponding regulatory regions (frag 1 and frag 3), or these regions with a mutated CArG-box (frags 1m and 3m), were tested with SOC1 and FUL homo- en heterodimers. The SOC1/FUL heterodimer could bind to both regulatory fragments, but this binding was reduced when the CArG-box was mutated in frag 1m and even completely abolished in frag 3m (**Fig. 3h**), providing additional evidence for the importance of the CArG-boxes for the binding of SOC1 and FUL. The SOC1 homodimer showed the same results, while the FUL homodimer did not show binding (**Supplementary Fig. 12**).

**Fig. 3.**
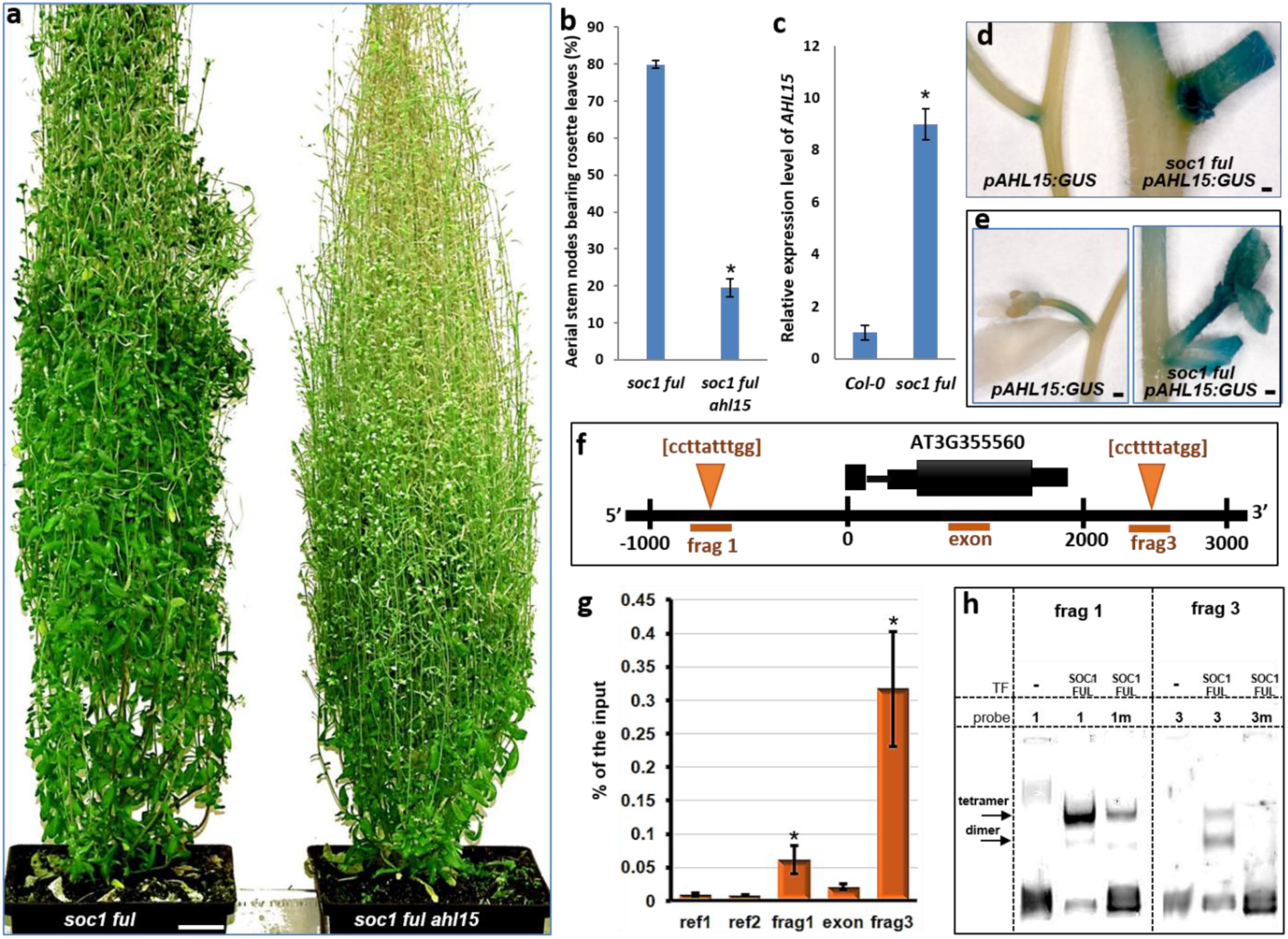
*AHL15* is essential for suppression of AM maturation in the Arabidopsis *soc1 ful* mutant. (**a**) A three-month-old *soc1 ful* double mutant plant with many aerial rosettes (left) and a *soc1 ful ahl15* triple mutant plant with a limited number of aerial rosettes (right), both grown in LD conditions. (**b**) Percentage of the aerial stem nodes bearing rosette leaves in three-month-old *soc1 ful* and *soc1 ful ahl15* plants. Asterisk indicates a significant difference (Student’s *t*-test, p < 0.001) and error bars represent standard error of the mean (n=10). (**c**) qPCR analysis of *AHL15* expression in secondary inflorescence nodes of wild-type (Col-0) and *soc1 ful* plants 2 weeks after flowering. Asterisk indicates a significant difference (Student’s *t*-test, p < 0.001) and error bars represent standard error of the mean (n=3). (**d** and **e**) *pAHL15:GUS* expression in an inflorescence node (**d**) or a secondary inflorescence (**e**) in wild-type (left) or *soc1 ful* mutant (right) background. Plants were of a comparable developmental stage. (**f**) *AHL15* gene model with canonical CArG boxes located in the upstream (frag 1) and downstream (frag 3) regions. (**g**) Graph showing ChIP-qPCR results from *FUL-GFP ap1 cal* secondary inflorescence nodes using anti-GFP antibody. The enrichment of the fragments was calculated as a percentage of the input sample. ref1/2, reference fragments (see Materials and Methods); Other fragments are indicated in the gene model. Error bars represent standard error of the mean (n=3). Significant differences from the control (Student’s *t*-test, p<0.01) are indicated with an asterisk. (**h**) Binding of SOC1 and FUL to regulatory regions near *AHL15*. Left panel: EMSA of promoter fragment 1 with a wild-type (1) or mutated (m1) CArG-box. Right panel: EMSA of downstream fragment 3 with a wild-type (3) or mutated (3m) CArG-box. Shifting of the probe, indicating binding, occurs either by a tetramer (upper band) or dimer (lower band). Size bar indicates 2 cm in **a** or 1 mm in **d** and **e**.

SOC1 and FUL are known as central floral integrators, as they integrate the different environmental and endogenous signaling pathways that influence flowering ^13,17,20^. They promote flowering through activation of the floral meristem genes *APETALA1* (*AP1*) and *LEAFY* (*LFY)* ^21,22^, and of genes involved in the biosynthesis of the plant hormone gibberellic acid (GA) ^23^. GA plays an important role in the promotion of flowering through activation of *SOC1* and the *SQUAMOSA PROMOTER BINDING PROTEIN-LIKE* (*SPL*) genes ^24,25^. Interestingly, *AHL15* and *AHL25* also control the GA biosynthesis through directly bind to the promoter of *GA3-oxidase1* (*GA3OX1*), which encodes an enzyme required for GA biosynthesis ^26^. We therefore investigated the relationship between GA biosynthesis and *AHL15* in the control of AM maturation. qPCR analysis showed that the expression of *GA3OX1, GA20OX1* and *GA20OX2*, genes, encoding rate limiting enzymes in the last steps of the GA biosynthetic pathway ^23,27,28^, was down-regulated in DEX-treated *35S:AHL15-GR* inflorescence nodes (**Fig. 4a**). In line with the down-regulation of GA biosynthesis, GA application to DEX-treated flowering *p35S:AHL15-GR* plants resulted in a remarkable repression of vegetative AM activity (**Fig. 4b**). In turn, treatment of flowering wild-type Arabidopsis plants by paclobutrazol, a potent inhibitor GA biosynthesis, prevented AM maturation, resulting in the aerial rosette leaf formation and enhanced longevity (**Fig. 4c**). Based on our findings, we postulate that *AHL15* acts downstream of SOC1 and FUL as central repressor of AM maturation, and that AHL15 prevents AM maturation in part by suppressing GA biosynthesis (**Fig. 4d**). Interestingly, the polycarpic behavior of *A. alpina* was shown to be based on age-dependent suppression of *AaSOC1* expression ^19^ and GA levels ^29^, and like in our model (**Fig. 4d**), *AHL* genes might also link these two regulatory pathways facilitating polycarpic growth in *A. alpina*.

**Fig. 4.**
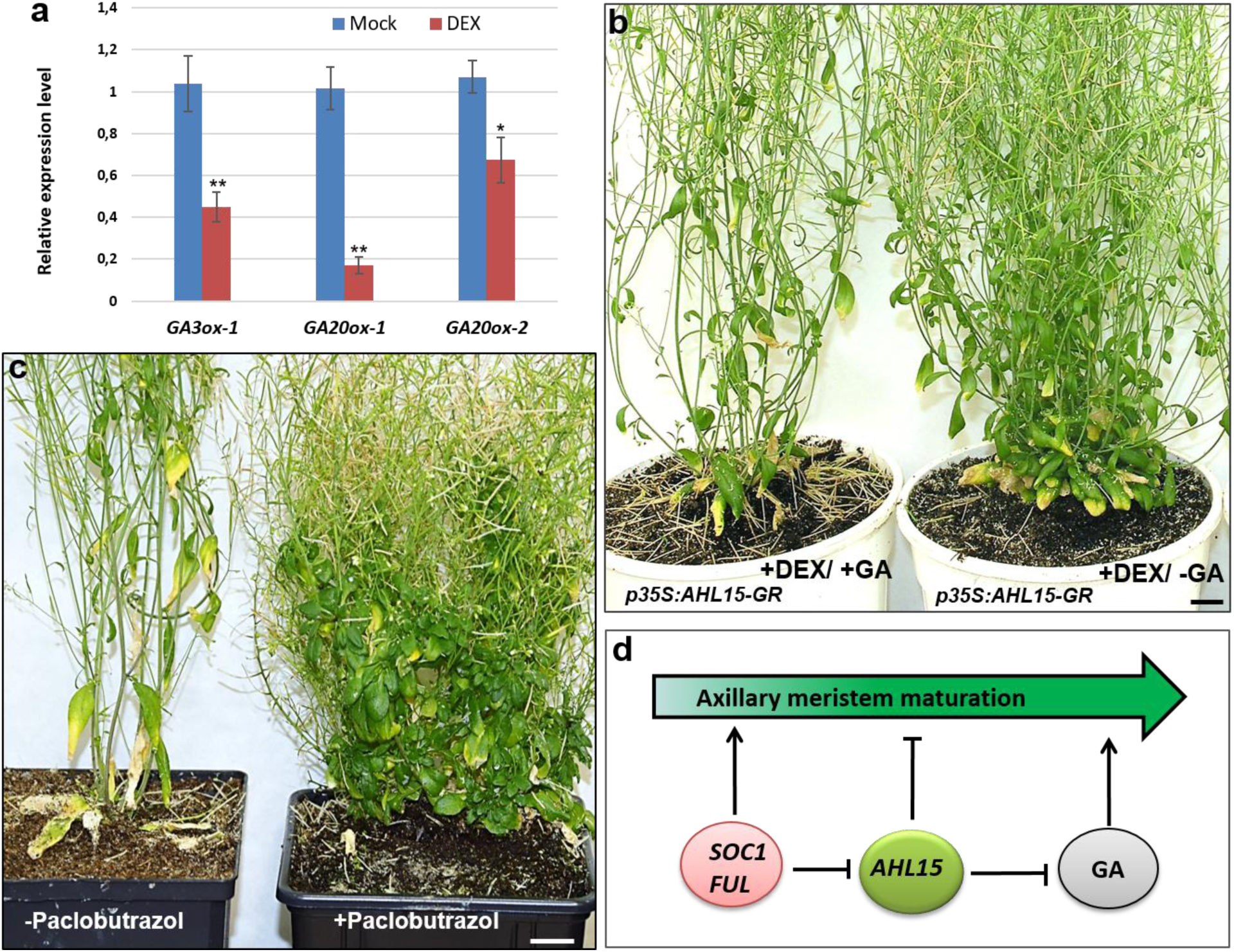
AHL15 delays AM maturation in part by suppressing gibberellic acid (GA) biosynthesis. (**a**) Relative expression level of the GA biosynthesis genes *GA3OX1, GA20OX1* and *GA20OX2* by qPCR analysis in the basal regions of 1-week-old *35S:AHL15-GR* inflorescences 1 day after spraying with water (mock) or with 20 μM DEX. Asterisks indicate a significant difference from mock-treated plants (Student’s *t*-test, * p<0.05, ** p<0.01). Error bars indicate standard error of the mean (n =3). (**b)** Shoot phenotype of 3-month-old *p35S:AHL15-GR* plants that were DEX sprayed as 35-day-old plants and subsequently sprayed 1 weak later with 10 μM GA_4_ (+GA) or with water (-GA). (**c)** Shoot phenotype of 3-month-old wild-type Arabidopsis plants that were sprayed 6 weeks earlier with water (-Paclobutrazol) or with 3μM paclobutrazol (+Paclobutrazol). Plants in **b** and **c** were grown in LD conditions and size bar indicates 2 cm. **(d)** Proposed model for the key role for *AHL15* in controlling AM maturation downstream of the flowering genes *SOC1* and *FUL* and upstream of GA biosynthesis. Blunted lines indicate repression, arrows indicate promotion.

The existence of both mono- and polycarpic species within many plant genera indicates that life history traits have changed frequently during evolution^5^. Clade-A *AHL* gene families can be found in both monocarpic and polycarpic plant species (**Supplementary Fig. 13a)** ^30^, and expression of the clade-A *AHL* family could therefore provide a mechanism by which a plant species attains a polycarpic growth habit. A comparison of the gene family size in representative species of 3 plant families did however not show significant gene deletion or duplication events linked to respectively the monocarpic or polycarpic growth habit (**Supplementary Fig. 13b)**. This suggests that a switch from monocarpic to polycarpic habit or vice versa is more likely to be mediated by a change in gene regulation.

To find indications for this, we compared the *AHL* gene family of Arabidopsis with that of its close polycarpic relative *Arabidopsis lyrata*^31^. The *A. thaliana* and *A. lyrata* genomes both encode 15 clade-A AHL proteins, among which the orthologous pairs can clearly be identified based on amino acid sequence identity (**Supplementary Fig. 14**). We hypothesized that the polycarpic habit of the monopodial *A. lyrata* is associated with enhanced clade-A *AHL* gene expression leading to repression of basal AM maturation during flowering (**Supplementary Fig. 15a**). Expression analysis of individual clade-A *AHL* genes in Arabidopsis showed that the expression of the majority of clade-A *AHL* genes, including *AHL15, AHL19*, and *AHL20*, was decreased in rosette nodes of *A. thaliana* flowering plants compared to 2 week old seedlings (**Supplementary Fig. 15b**). In contrast, the expression of 5 members of the *AHL* gene family (*AHL15, AHL17, AHL19, AHL20* and *AHL27)* was significantly higher in rosette nodes of flowering *A. lyrata* plants compared to seedlings (**Supplementary Fig. 15c**). These data are in line with our hypothesis, and suggest that the different life-history strategies in *A. thaliana* and *A. lyrata* might be determined by differential regulation of *AHL* genes in AMs.

In conclusion, our data provide evidence for a novel role for *AHL15* and most likely other clade-A *AHL* genes in enhancing plant longevity by suppressing AM maturation. Although the exact mode of action of AHL proteins is largely unknown, they are characterized as DNA-binding proteins, and like AT-hook proteins in animals they seem to act through chromatin remodeling ^11,32^. It has been shown that AHL22 represses *FLOWERING LOCUS T* (*FT*) expression by binding to the *FT* promoter where it possibly modulates the epigenetic signature around its binding region^12^. Detailed studies on the chromatin configuration by approaches such as chromosome conformation capture technologies ^33^ should provide more insight into the mode of action of these plant-specific AT-Hook motif proteins. One of the objectives of our future research will be to unravel the molecular mechanisms by which these proteins influence plant development.

## ACKNOWLEDGEMENTS

We thank Kim Boutilier and Thomas Greb for critical comments on the manuscript. O.K. was financially supported by a grant from the Iran Ministry of Science, Research and Technology (MSRT 89100156), and by subsidies from Generade and the Building Blocks of Life research programme (project number 737.016.013 to R.O.), which is partly financed by the Netherlands Organisation for Scientific Research (NWO). M.K. was financially supported by a fellowship of the Institute of Biotechnology & Genetic Engineering (IBGE) at the Agricultural University of Peshawar with financial support by the Higher Education Commission (HEC) of Pakistan. R.R.H. was financially supported by the KU Leuven Research Fund. M.B. was supported by grant ALWOP.199 from NWO. P.M. and M.C. were supported in part by an Innovation Subsidy Collaborative Projects (IS054064) from the Dutch Ministry of Economic Affairs.

## AUTHOR CONTRIBUTIONS

O.K. and R.O. conceived and supervised the project; All authors designed experiments and analyzed and interpreted results; O.K. performed the majority of the Arabidopsis experiments; A.R., M.B., P.M., and M.C. contributed to the Arabidopsis experiments; M.K. generated and analyzed the tobacco lines; R.R.H. and V.N. analysed the *AHL* gene families in different mono- and polycarpic plant species, O.K. and R.O. wrote the manuscript; All authors read and commented on versions of the manuscript.

## METHODS

### Plant material, growth conditions and phenotyping

All Arabidopsis mutant- and transgenic lines used in this study are in the Columbia (Col-0) background. The *ahl15* (SALK_040729) T-DNA insertion mutant and the previously described *soc1-6 ful-7* double mutant^34^ were obtained from the Nottingham Arabidopsis Stock Centre (NASC). Seeds were snow soil and grown at 21°C, 65% relative humidity and a 16 hours (long day: LD) or 8 hours (short day: SD) photoperiod. To score for phenotypes such as longevity, Col-0 wild-type, mutant or transgenic plants were transferred to larger pots about 3 weeks after flowering. *Nicotiana tabacum* cv SR1 Petit Havana (tobacco) plants were grown in medium-sized pots at 25 C°, 70% relative humidity and a 16 hours photoperiod. For dexamethasone (DEX, Sigma-Aldrich) treatment, Arabidopsis and tobacco plants were sprayed with 20 and 30 µM DEX, respectively. To test the effect of GA on AHL15-GR activation by DEX treatment, 35-day-old flowering *p35S:AHL15-GR* plants were first sprayed with 20 µM DEX, followed 1 week later by spraying with 10 μM GA4 (Sigma-Aldrich). The production of rosette leaves, cauline leaves, flowers or fruits per rosette or aerial AM was determined by dividing the total number of leaves or fruits produced by the number of active rosette or aerial AMs per plant. For the flowering time the number of rosette leaves produced by the SAM were counted upon bolting.

### Plasmid construction and transgenic Arabidopsis lines

To generate the different *promoter:AHL15* gene fusions, the complete *AHL15* (AT3G55560) genomic fragment from ATG to stop codon was amplified from genomic DNA of Arabidopsis ecotype Columbia (Col-0) using PCR primers Gateway-AHL15-F and -R (Supplementary Table 1). The resulting fragment was inserted into pDONR207 via a BP reaction. LR reactions were carried out to fuse the *AHL15* downstream of the *35S* promotor in destination vector pMDC32 ^35^. Subsequently, the *35S* promoter was excised with *Kpn*I and *Sph*I and replaced by the Gateway cassette (*ccdB* flanked by *attP* sequences) amplified from pMDC164^35^ by the *Kpn*I and *Sph*I flanked primers (Supplementary Table 1), resulting in plasmid pGW-AHL15. To generate the constructs *pFD:AHL15, pMYB85:AHL15, pMYB103:AHL15* and *pAHL15:AHL15*, 3 kb regions upstream of the ATG initiation codon of the genes *FD* (AT4G35900), *MYB85* (AT4G22680), *MYB103* (AT1G63910) and *AHL15* were amplified from ecotype Columbia (Col-0) genomic DNA using the forward (F) and reverse (R) PCR primers indicated in Supplementary Table 1. The resulting fragments were first inserted into pDONR207 by BP reaction, and subsequently cloned upstream of the *AHL15* genomic fragment in destination vector pGW-AHL15 by LR reaction. To generate the *pAHL15:GUS, pMYB85:GUS* and *pMYB103:GUS* reporter constructs, the corresponding promotor fragments were cloned upstream of GUS in destination vector pMDC164 by LR reaction. To generate the *pAHL15:AHL15-ΔG* construct, a synthetic *Kpn*I-*Spe*I fragment containing the *AHL15* coding region lacking the sequence encoding the Gly-Arg-Phe-Glu-Ile-Leu amino acids in the C-terminal region (BaseClear) was used to replace the corresponding coding region in the *pAHL15:AHL15* construct. To construct *35S::AHL15-GR*, a synthetic *Pst*I-*Xho*I fragment containing the *AHL15-GR* fusion (Shine Gene Molecular Biotech, Inc: Supplementary File. 1) was used to replace the BBM-GR fragment in binary vector pSRS031^36^. To generate the other overexpression constructs, the full-length cDNA clones of *AHL19* (AT3G04570), *AHL20* (AT4G14465), *AHL27* (AT1G20900) and *AHL29* (AT1G76500) from Arabidopsis Col-0, *AC129090* from *Medicago trunculata cv* Jemalong (*MtAHL15*), and *Bo-Hook1* (AM057906) from *Brassica oleracea* var *alboglabra* (*BoAHL15*) were used to amplify the open reading frames using primers indicated in Supplementary Table 1. The resulting fragments were cloned into plasmid pJET1/blunt (GeneJET™ PCR Cloning Kit, #K1221), and subsequently transferred as *Not*I fragments to binary vector pGPTV 35S-FLAG ^37^. All binary vectors were introduced into *Agrobacterium tumefaciens* strain AGL1 by electroporation ^38^ and Arabidopsis Col-0 and *ahl15* plants were transformed using the floral dip method ^39^.

### Tobacco transformation

Round leaf discs were prepared from the lamina of 3rd and 4th leaves of 1-month-old soil grown tobacco plants. The leaf discs were surface sterilized by three washes with sterile water followed by incubation in 10 % glorix for 20 minutes^40^, and then again 4 to 5 washes with sterile water. The surface sterilized leaf discs were syringe infiltrated with an overnight acetosyringone (AS)-induced culture of *Agrobacterium tumefaciens* strain AGL1 containing binary vector pSRS031 (grown to OD_600_= 0.6 in the presence of 100 µM AS) carrying the *35S::AHL15-GR* construct and co-cultivated for 3 days in the dark on co-cultivation medium (CCM), consisting of full strength MS medium ^41^ with 3% (w/v) sucrose (pH 5.8) solidified with 0.8 % (w/v) Diachin agar and supplemented with 2mg/l BAP, 0.2mg/l NAA and 40mg/l AS. After co-cultivation, the explants were transferred to CCM supplemented with 15mg/l phosphinothricin (ppt) for selection and 500mg/l cefotaxime to kill *Agrobacterium*. Regeneration was carried out at 24C° and 16 hours photoperiod. The regenerated transgenic shoots were rooted in big jars containing 100 ml hormone free MS medium with 15mg/l ppt and 500 mg/l cefotaxime. The rooted transgenic plants were transferred to soil and grown in a growth room at 25 °C, 75% relative humidity and a 16 h photoperiod. All the transgenic plants were checked for the presence of the T-DNA insert by PCR, using genomic DNA extracted from leaf tissues by the CTAB method ^42^.

### Histolochemical staining and microscopy

Histochemical β-glucuronidase (GUS) staining of transgenic lines expressing GUS was performed as described previously ^43^. Tissues were stained for 4 hours at 37°C, followed by chlorophyll extraction and rehydration by incubation for 10 minutes in a graded ethanol series (75, 50, and 25 %). GUS stained tissues were observed and photographed using a LEICA MZ12 microscope (Switzerland) equipped with a LEICA DC500 camera.

### Quantitative real-time PCR (qPCR) analysis

RNA isolation was performed using a NucleoSpin® RNA Plant kit (MACHEREY-NAGEL). For qPCR analysis, 1 µg of total RNA was used for cDNA synthesis with the iScript™ cDNA Synthesis Kit (BioRad). PCR was performed using the SYBR-Green PCR Master mix (SYBR® Premix Ex Taq™, Takara) and a CFX96 thermal cycler (BioRad). The Pfaffl method was used to determine relative expression levels ^44^. Expression was normalized using the *β-TUBULIN-6* and *EF1-ALPHA* genes. Three biological replicates were performed, with three technical replicates each. The primers used are described in Supplementary Table 2.

### ChIP-qPCR experiment

For the ChIP-qPCR analysis, three independent samples were harvested from secondary inflorescence nodes of *pFUL:FUL-GFP ful* plants and processed as described in ^45,46^. Primer sequences used for the ChIP-qPCR are detailed in Supplementary Table 2.

### EMSA experiment

The EMSA was performed as described in Bemer et al. 2017 ^47^. The sequences of the probes are detailed in Supplementary Table 2.

### Clade-A *AHL* gene family data retrieval

The nucleotide and amino acid sequences for Clade-A *AHLs* in *A. thaliana* (*AtAHLs*) were retrieved by Biomart from Ensembl Plants (plants.ensembl.org/index.html). For our study we selected 15 additional species from 3 major plant families, i.e. Brassicaceae, Solanaceae and Fabaceae. Initially more species were included, but some were excluded from the analysis (e.g. *Arabis alpina*) for reasons described below. The amino acid sequences of *A. thaliana, A. lyrata, Brassica oleracea, Brassica rapa, Solanum lycopersicum, Solanum tuberosum, Medicago truncatula* and *Glycine max* were downloaded from Ensembl Plants (ftp://ftp.ensemblgenomes.org). The genomes of *Nicotiana tabacum, Capsicum annuum, Brassica napus* were downloaded from NCBI Genome (ftp://ftp.ncbi.nih.gov/genomes/) and the genomes of *Phaseolus vulgaris, Capsella rubella, Capsella grandiflora, Boechera stricta* and *Eutrema salsugineum* were downloaded from Phytozome (https://phytozome.jgi.doe.gov/pz/portal.html). The above species were selected such that at least one monocarpic and one polycarpic species was represented within each of the 3 families.

### Building of profile HMMs and hmmer searches

The whole protein sequences of the Arabidopsis Clade-A AHLs were used as a query and BLASTP ^48^ with an e-value set at 0.001 was used to search for AHLs in the other plant genomes. Only BLASTP hits >70% coverage and 70% sequence identity with intact single AT-Hook motif and PPC domain were used for building profile Hidden Markov Models (HMMs) for Clade-A AHLs. We used MAFFT software ^49^ with FF-NS-i algorithm for construction of seed alignments. The alignments were manually inspected to remove any doubtful sequences. To increase the specificity of the search, columns with many gaps or low conservation were excluded using the trimAl software ^50^. We applied a strict non-gap percentage threshold of 80% or similarity score lower than 0.001 such that at least 30% of the columns were conserved. At this point several species were excluded (e.g. *Arabis alpina*), because of extensive gaps in the sequence alignment. Profile HMMs were built from the Multiple Sequence Alignment (MSA) aligned fasta files using hmmbuild and subsequent searches against the remaining 16 genomes was carried out using hmmsearch from the HMMER 3.1b1 package ^51^. AHL proteins in plants consist of two closely resembling clades; Clade-A and Clade-B. AHL sequences were classified into Clade-A family based on a comparison with Clade-B AHL sequences where a hit with lower e-value for either Clade-A or Clade-B would correctly place the sequence in the corresponding clade (e.g. low e-value for Clade A would place the sequence in Clade-A and vice-versa.)

### Phylogenetic reconstruction and reconciliation

Phylogenetic analysis was carried out using both Maximum Likelihood (ML) with PhyML^52^ and Bayesian Inference implementing the Markov Chain Monte Carlo (MCMC) algorithm with MrBayes ^53^. For Bayesian inference, we specified the number of substitution types (nst) equal to 6 and the rate variation (rates) as invgamma. Invgamma states that a proportion of the sites are invariable while the rate for the remaining sites are drawn from a gamma distribution. These settings are equivalent to the GTR + I + gamma model. Two independent analysis (nruns=2) of 4 chains (3 heated and one cold) were run simultaneously for at least 10 million generations, sampling every 1000 generations. Burn-in was set as 25%. For Clade-A AHLs the simulations were run for 10 million generations, sampling every 1000 generations and convergence was reached at 0.016. For ML analysis, we used the default amino acid substitution model LG and the number of bootstrap replicates was specified as 100.

Tree resolving, rearrangement, and reconciliation was carried out using NOTUNG software ^54^. NOTUNG uses duplication/loss parsimony to fit a gene (protein) tree to a species tree. The species tree was obtained using PhyloT (http://phylot.biobyte.de/index.html) which generates phylogenetic trees based on the NCBI taxonomy. Tree editing/manipulations were performed using the R package; APE ^55^ and GEIGER ^56^. We applied a strict threshold for rearrangement of 90%. After the rearrangement, we performed the reconciliation of the gene (protein) tree with the species tree.

### Reconstruction of evolutionary scenario using Dollo parsimony method

Dollo parsimony principles are commonly exploited for two-state character traits. To classify branches as either gene-losses or gene-gains, we used Dollo parsimony method, which allows for an unambiguous reconstruction of ancestral character states as it is based on the assumption that a complex character that has been lost during evolution of a particular lineage cannot be regained.

