## Supplementary Figures for "A suppressor of axillary meristem maturation promotes longevity in flowering plants"

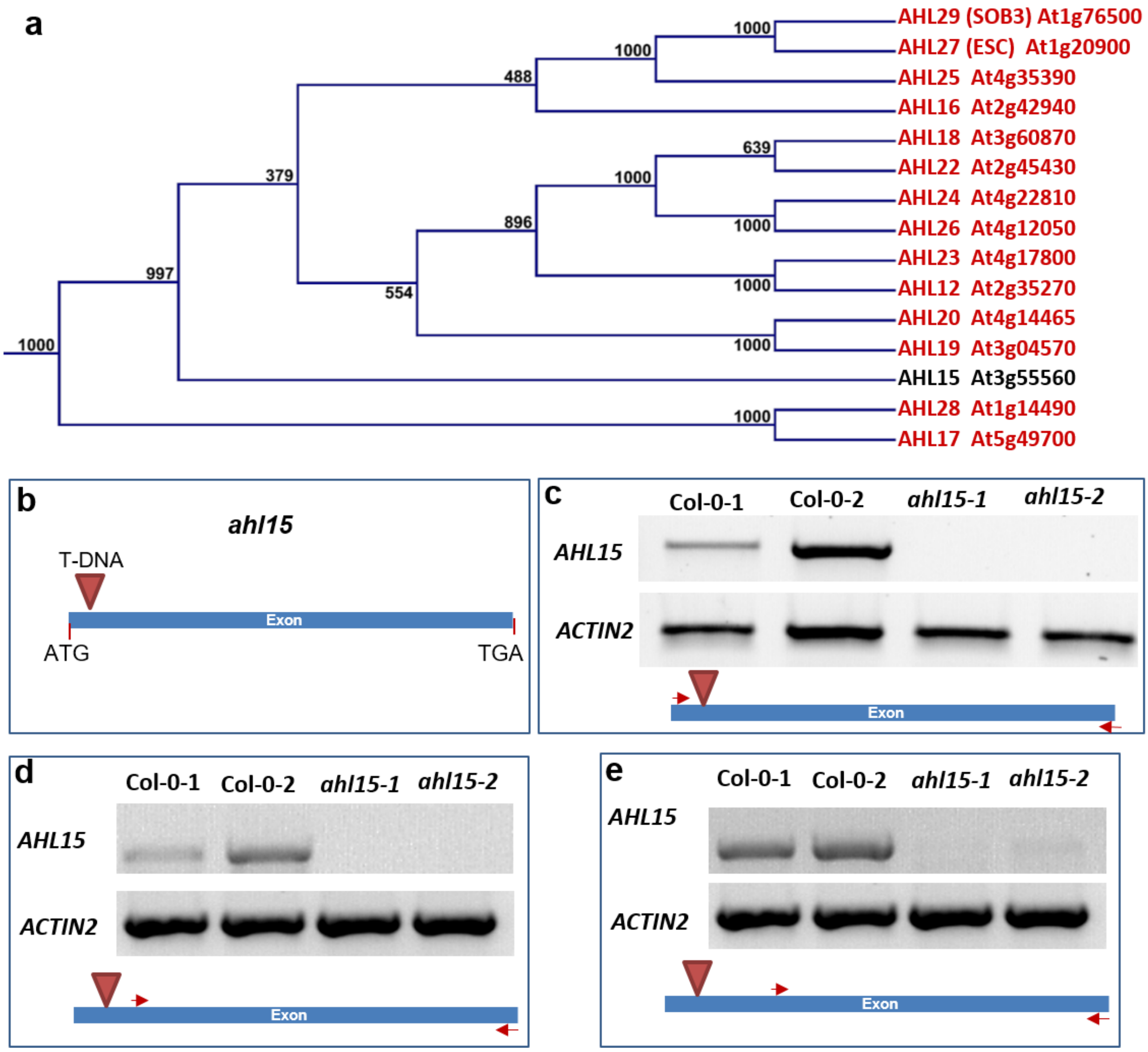


**Supplementary Figure 1 | The Arabidopsis clade-A AHL protein and gene family.** (**a**) A phylogenetic tree of part of the Arabidopsis AHL protein family showing all clade-A AHL proteins containing a single At-Hook domain, including AHL15. (**b**) Location of the T-DNA insertion in the intronless *AHL15* gene, resulting in the *ahl15* loss-of-function mutant allele used for this research. (**c-e**) Duplicate RT-PCR detection of full-length *AHL15* coding sequence (CDS) (**c**), from 64bp downstream of the ATG until the end of the CDS (**d**) and from 197bp downstream of the ATG until end of the CDS (**e**) in wild-type Arabidopsis (Col-0) but not in the *ahl15* mutant. The *ACTIN2* was used as a positive control. The primers used for RT-PCR are described in Supplementary Table 2.

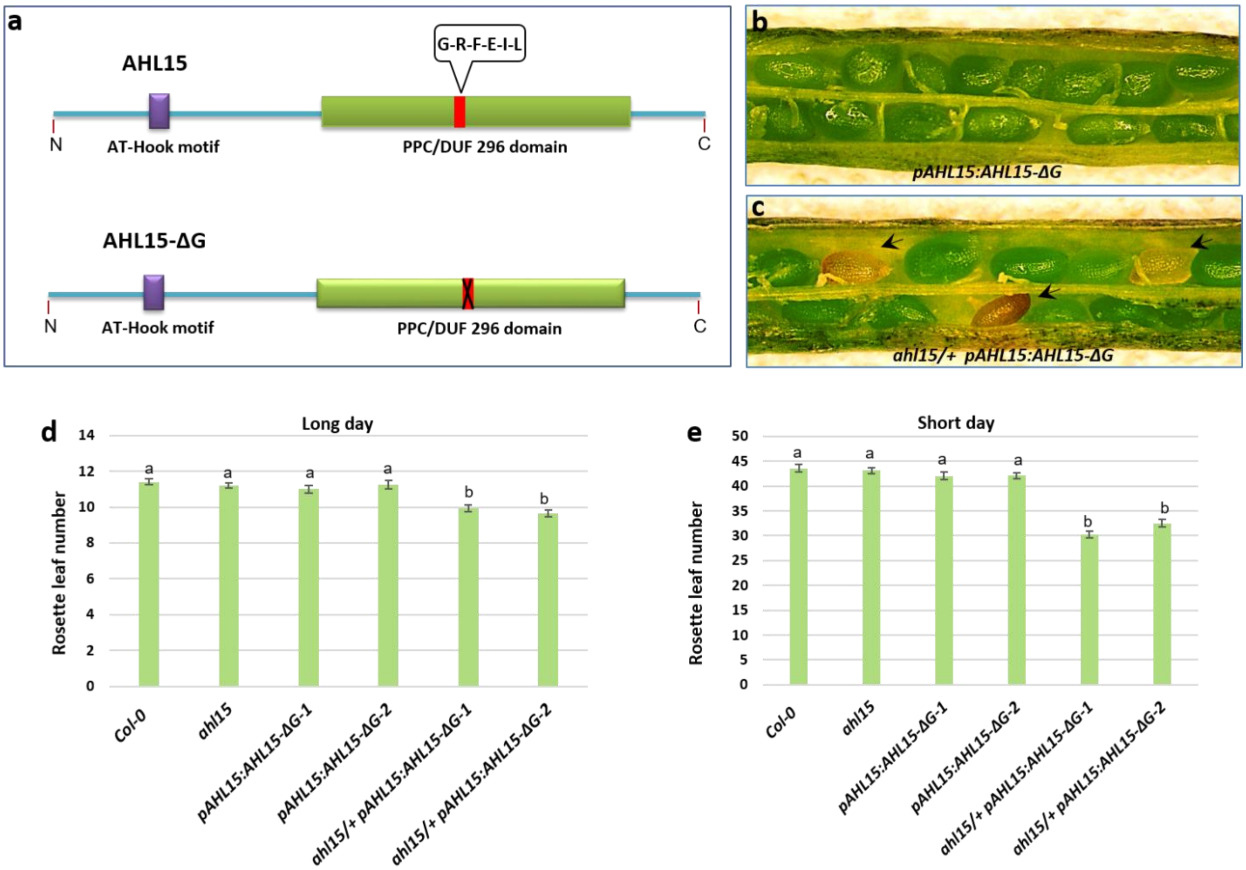


**Supplementary Figure 2 | Expression of a dominant negative AHL15-ΔG mutant protein in the Arabidopsis *ahl15* mutant background causes early flowering and impairs seed development.** (**a**) The schematic domain structure of AHL15 and the dominant negative AHL15-ΔG version, in which six-conserved amino-acids (Gly-Arg-Phe-Glu-Ile-Leu, red box) are deleted from the C-terminal PPC domain. **(b)** Wild-type seed development in *pAHL15:AHL15-ΔG* siliques. **(c)** Aberrant seed development (arrowheads) in *ahl15*/+ *pAHL15:AHL15-ΔG* siliques (observed in 3 independent *pAHL15:AHL15-ΔG* lines crossed with the *ahl15* mutant). (**d, e**) The number of rosette leaves produced by the SAM in wild-type, *ahl15*, *pAHL15:AHL15-ΔG* and *ahl15/+ pAHL15:AHL15-ΔG* plants (n=15) plants grown in long day (LD, **d**) or short day (SD, **e**) conditions. Two independent transgenic lines (1 and 2) were used in each experiment. Error bars represent standard eror of the mean. Letters (a and b)

indicate statistically significant differences, as determined by a one-way ANOVA with a Tukey's HSD post hoc test ($p < 0.01$).

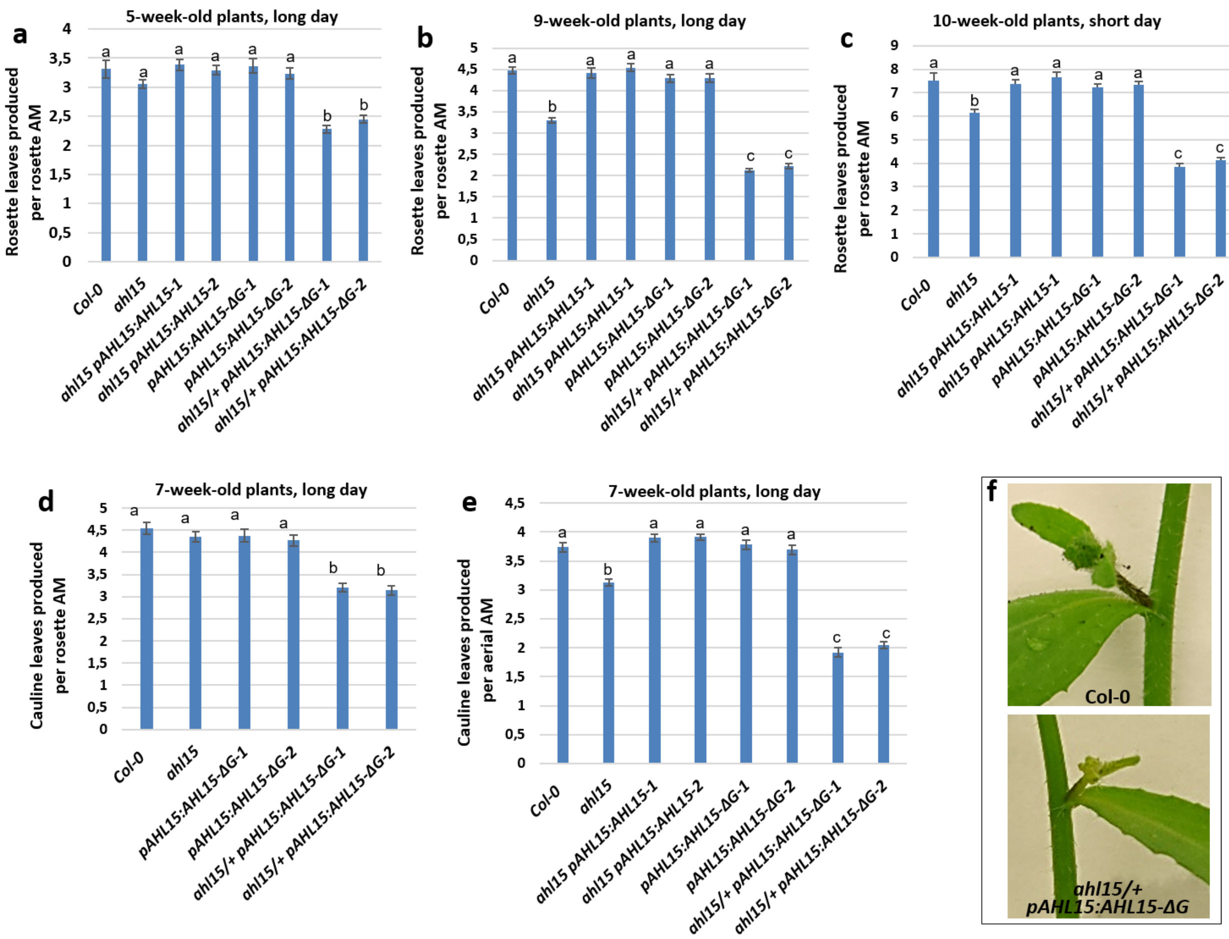


**Supplementary Figure 3 | *AHL15* represses AM maturation in Arabidopsis**. (**a, b**) The number of rosette leaves produced per rosette AM of wild-type, *ahl15*, *ahl15 pAHL15:AHL15*, *pAHL15:AHL15-ΔG* and *ahl15/+ pAHL15:AHL15-ΔG* plants (n=15) 5 (**a**), 9 (**b**), or 10 weeks (**c**) after germination in long day (LD, **a**,**b**) or short day (SD, **c**) conditions. (**d, e**) The number of cauline leaves produced by rosette AMs (**d**) or by aerial AMs (**e**) of 7-week-old wild-type, *ahl15*, *ahl15 pAHL15:AHL15*, *pAHL15:AHL15-ΔG* and *ahl15/+ pAHL15:AHL15-ΔG* plants (n=15). (**f**) A lateral inflorescence with cauline leaves formed on the first inflorescence node of a wild-type (**top**) or *ahl15/+ pAHL15:AHL15-ΔG* plant (**bottom**). Error bars in **a-e** represent the standard error of the mean. Letters (a, b, c) indicate statistically significant differences, as determined by a one-way ANOVA with a Tukey's HSD post hoc test ($p < 0.01$).

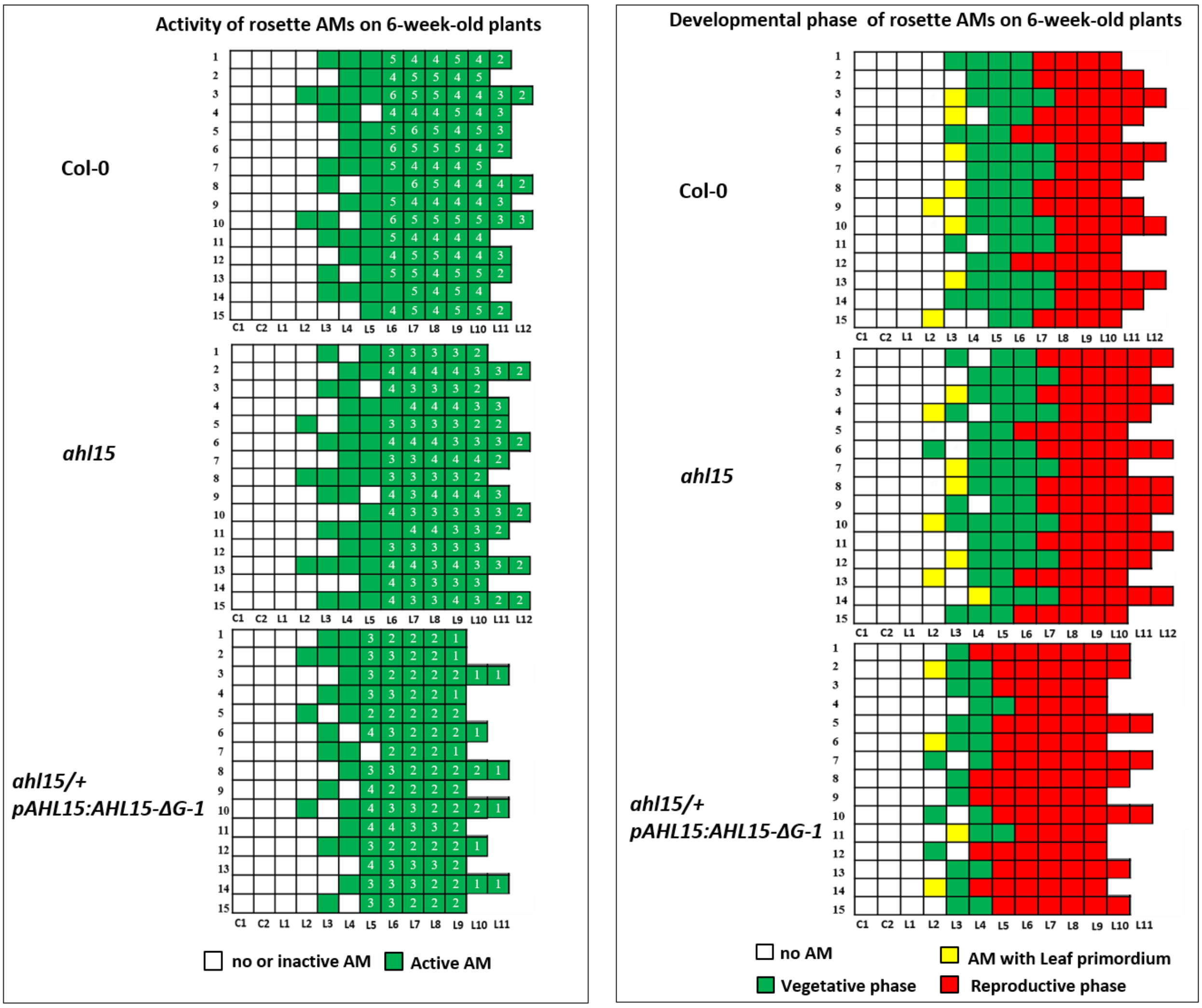


**Supplementary Figure 4 | Arabidopsis *AHL* genes enhance the vegetative activity and suppress the floral transition of rosette AMs.** (**a**) Schematic representation of the vegetative activity of rosette AMs located of six-week-old wild-type, *ahl15* and *ahl15/+ pAHL15:AHL15-ΔG-1* plants. Each row represents a single plant, and each square represents an individual AM in a cotyledon axil (C1 and C2) or in a rosette leaf axils (L1 to L12). The numbers within a square represent the number of rosette leaves produced by a rosette AM. A green square indicates a leaf axil with an active AM, as indicated by bud growth or leaf development, and a white square indicates a leaf axil without an (active) AM. (**b**) Developmental phase of the rosette AMs of six-week-old wild-type, *ahl15* and *ahl15/+ pAHL15:AHL15-ΔG-1* plants. White, yellow, green or red squares indicate axils without (active) AM, or rosette AMs with at least one visible leaf primordium, producing rosette leaves (vegetative) or producing cauline leaves or flowers (reproductive), respectively. Plants in **a** and **b** were grown in LD conditions.

**Supplementary Figure 5 | Repression of AM maturation by *AHL15* leads to increased**

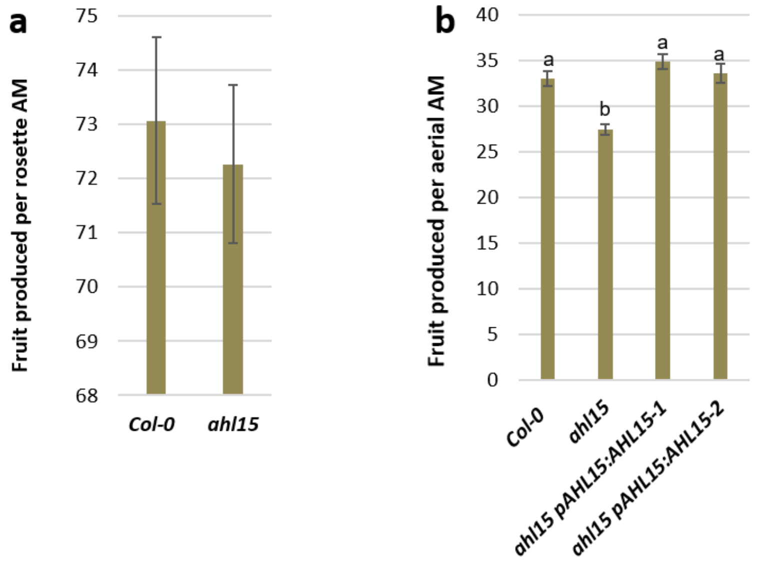


**flower and fruit production in Arabidopsis.** (**a**) The number of fruits produced by the rosette AMs of 65-day-old wild-type or *ahl15* plants (n=15, Student's t-test, $p < 0.01$). (**b**) The number of fruits produced by the aerial AMs of 65-day-old wild-type, *ahl15* or *ahl15 pAHL15:AHL15* plants. Error bars in **a** and **b** represent the standard error of the mean. Letters (a, b, c) in **b** indicate statistically significant differences, as determined by a one-way ANOVA with a Tukey's HSD post hoc test ($p < 0.01$).

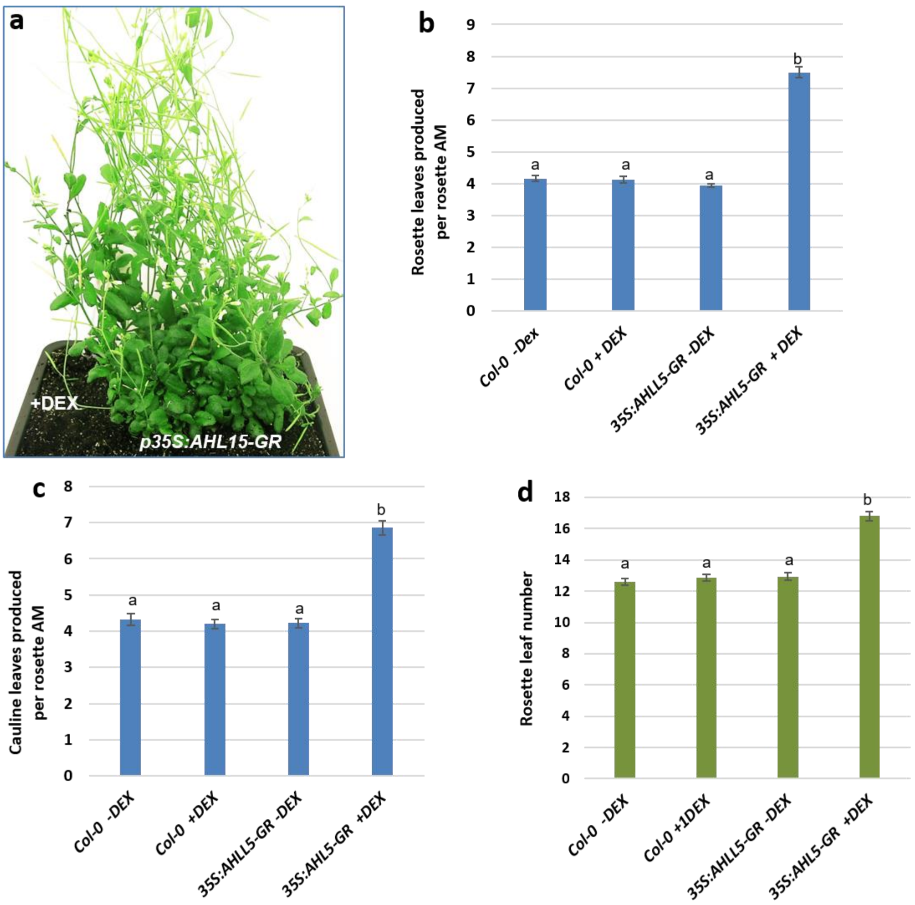


**Supplementary Figure 6 | *AHL15* overexpression delays floral transition of the SAM and represses AM maturation.** (**a**) Shoot phenotype of a flowering 7-week-old *35S:AHL15-GR* plant that was DEX-treated upon bolting (approx. 5 weeks old). (**b**) Number of rosette leaves produced by rosette AMs of 7-week-old mock-treated wild-type, DEX-treated wild-type, mock-treated *35S:AHL15-GR* and DEX-treated *35S:AHL15-GR* plants (n=15). (**c**) Number of cauline leaves produced by the rosette AMs of 7-week-old mock-treated wild-type, DEX-treated wild-type, mock-treated *35S:AHL15-GR* and DEX-treated *35S:AHL15-GR* plants (n=15). Plants in **b** and **c** were treated upon bolting (5 weeks old) and scored 2 weeks later. (**d**) The number of rosette leaves produced by the SAM in mock-treated wild-type, DEX-treated wild-type, mock-treated *35S:AHL15-GR* and DEX-treated *35S:AHL15-GR* plants (n=15). Non-flowering (3-week-old) plants were treated and the SAM-produced rosette leaves were counted after bolting. Error bars in **b, c and d** represent the standard error of the mean (SEM). Letters (a, b, c) indicate statistically significant differences, as determined by a one-way ANOVA with a Tukey's HSD post hoc test ($p < 0.01$). Plants were grown in LD conditions.

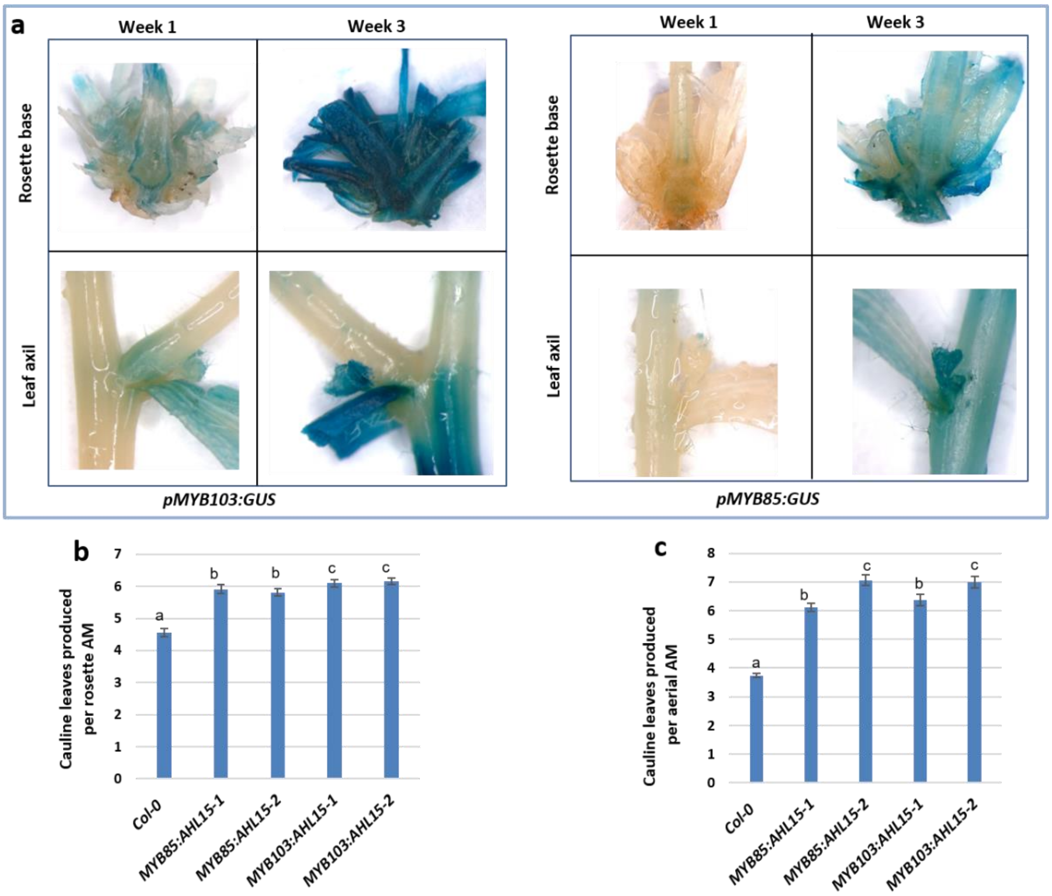


**Supplementary Figure 7 | *AHL15* overexpression in the rosette base and leaf axils delays AM maturation in Arabidopsis.** (**a**) Expression of *pMYB58:GUS* and *pMYB103:GUS* reporters in the rosette base (top) or leaf axils (bottom) of Arabidopsis plants respectively one or three weeks after flowering, as monitored by histochemical GUS staining. (**b, c**) The number of cauline leaves produced by rosette AMs (**b**) or aerial AMs (**c**) of 60-day-old (**b**) or 65-day-old (**c**) wild-type, *pMYB85:AHL15* or *pMYB103:AHL15* plants grown in LD conditions (n=15). Error bars represent the standard error of the mean. Letters (a, b, c) indicate statistically significant differences, as determined by a one-way ANOVA with a Tukey's HSD post hoc test ($p < 0.01$).

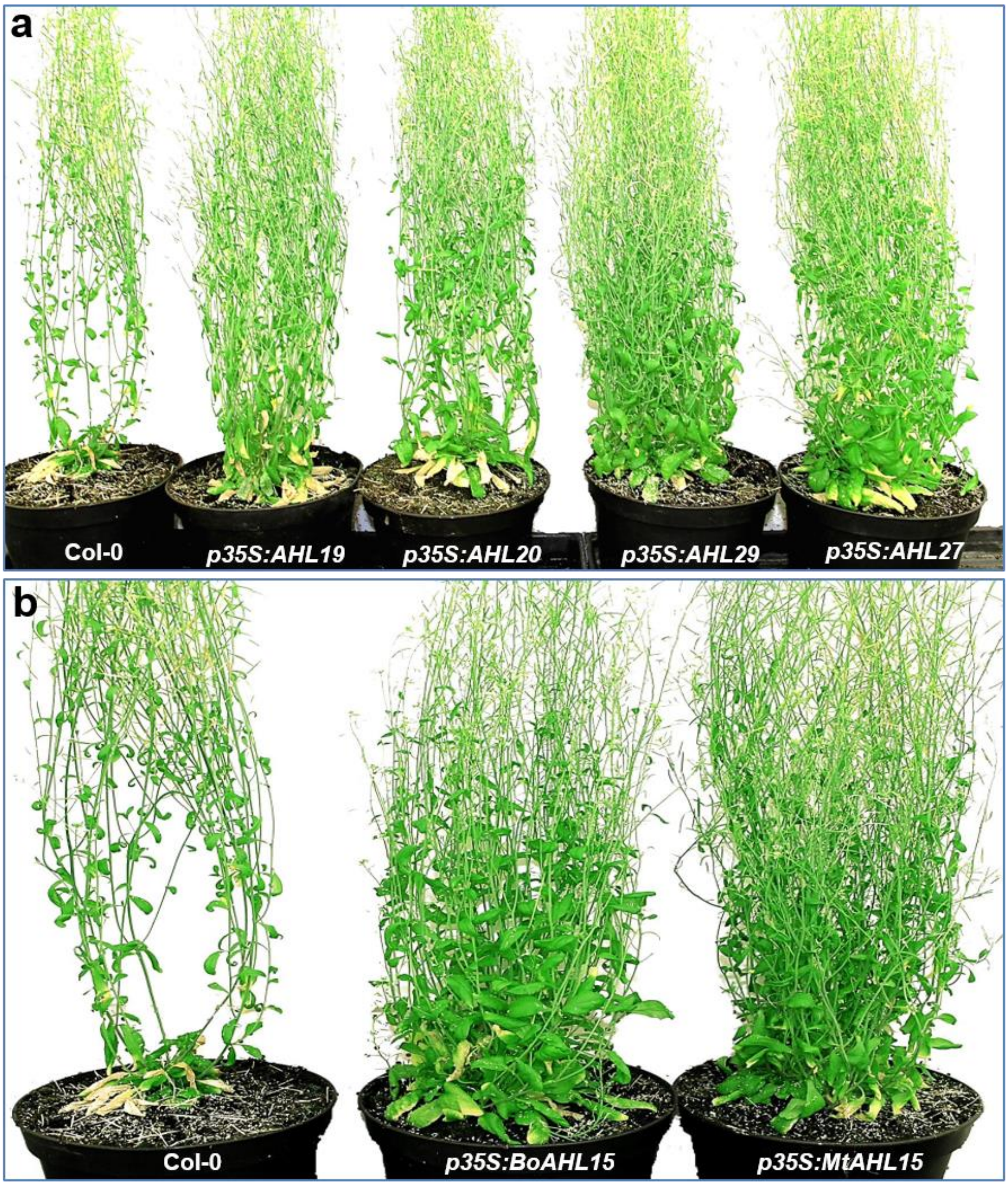


**Supplementary Figure 8 | Overexpression of Arabidopsis *AHL15* paralogs or putative orthologs represses AM maturation in Arabidopsis**. (**a** and **b**) Wild-type (Col-0) or transgenic Arabidopsis plants overexpressing Arabidopsis *AHL19*, *AHL20*, *AHL27* and *AHL29* (**a**), or the putative *AHL15* orthologs from *Brassica oleracea* (*BoAHL15*) or *Medicago trunculata* (*MtAHL15*) (**b**). Plants were grown in LD conditions. For presentation purposes, the original background of the images was replaced by a homogeneous white background.

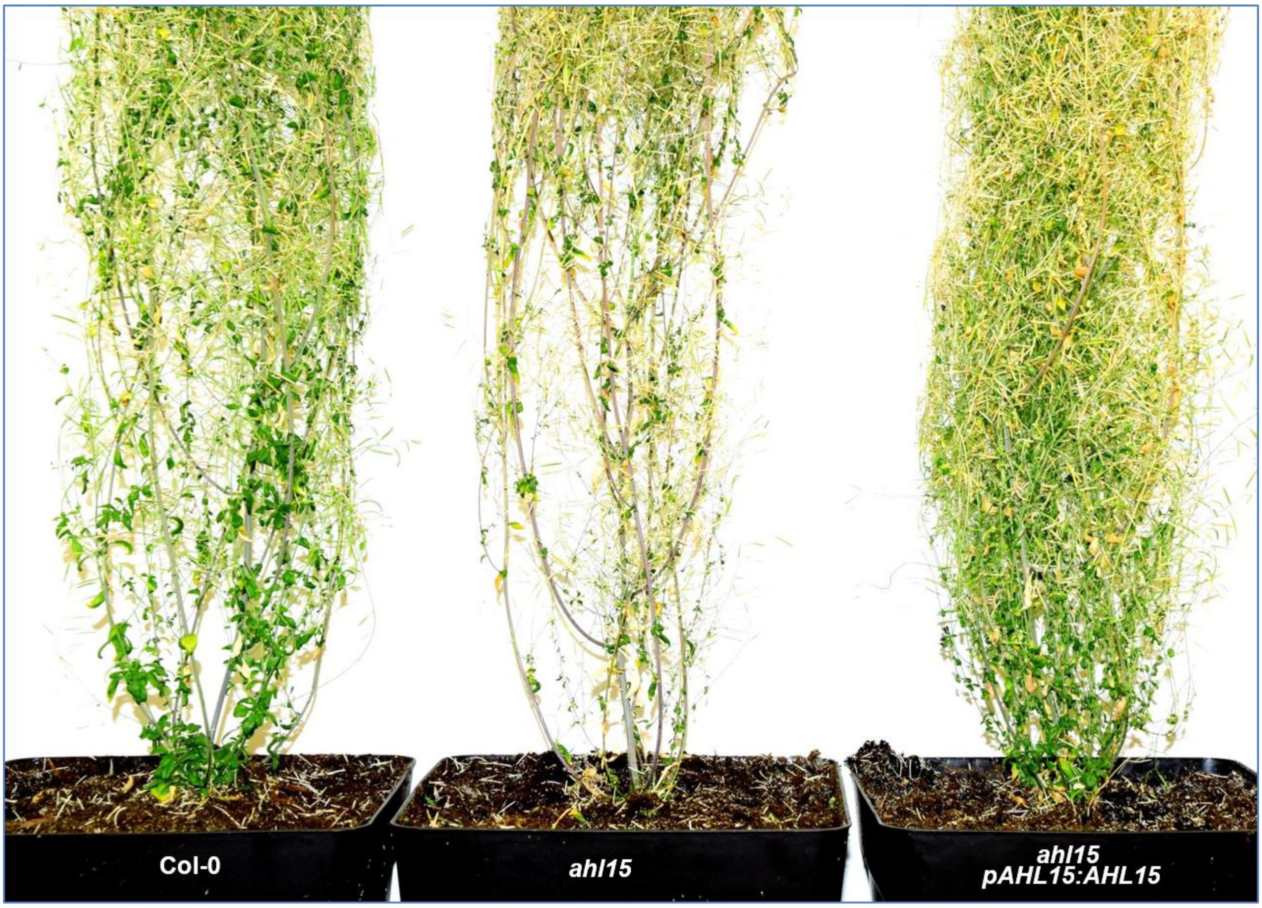


**Supplementary Figure 9 | *AHL15* enhances the longevity of short day-grown Arabidopsis plants.** (**c**) Phenotype of 5-month-old wild-type (Col-0, left), *ahl15* (middle) and *ahl15 pAHL15:AHL15* (right) plants. The plants were grown in SD conditions. For presentation purposes, the original background of the image in was replaced by a homogeneous white background.

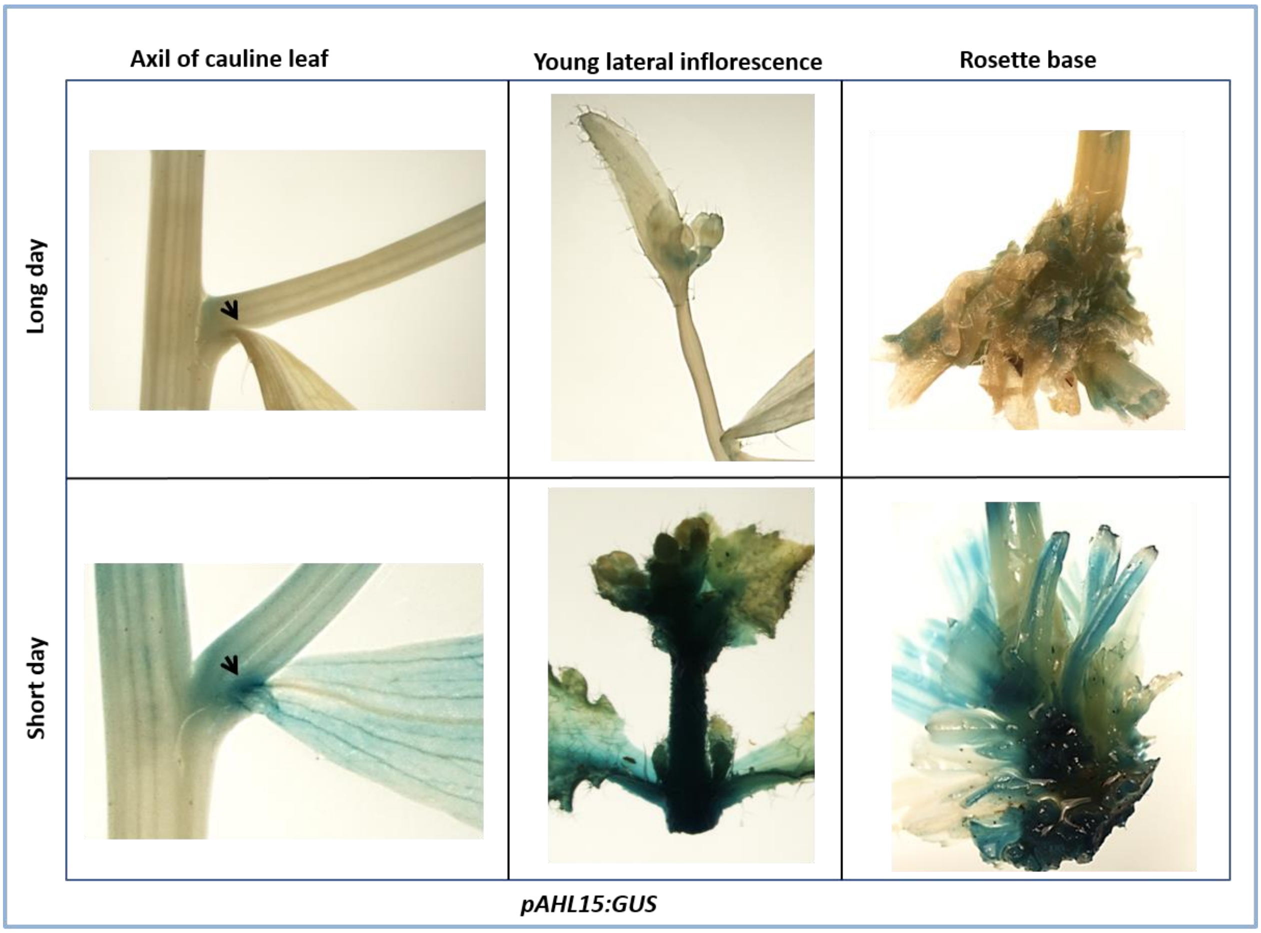


**Supplementary Figure 10 | *AHL15* expression is day length sensitive.** Expression of the *pAHL15:GUS* reporter in the axil of a cauline leaf (**lef**t, **arrowheads**), Young lateral inflorescence (**middle**) and rosette base (**right**) of a 9-week-old plant grown in LD conditions (**top**) or a 4-month-old plant grown under SD conditions (**bottom**).

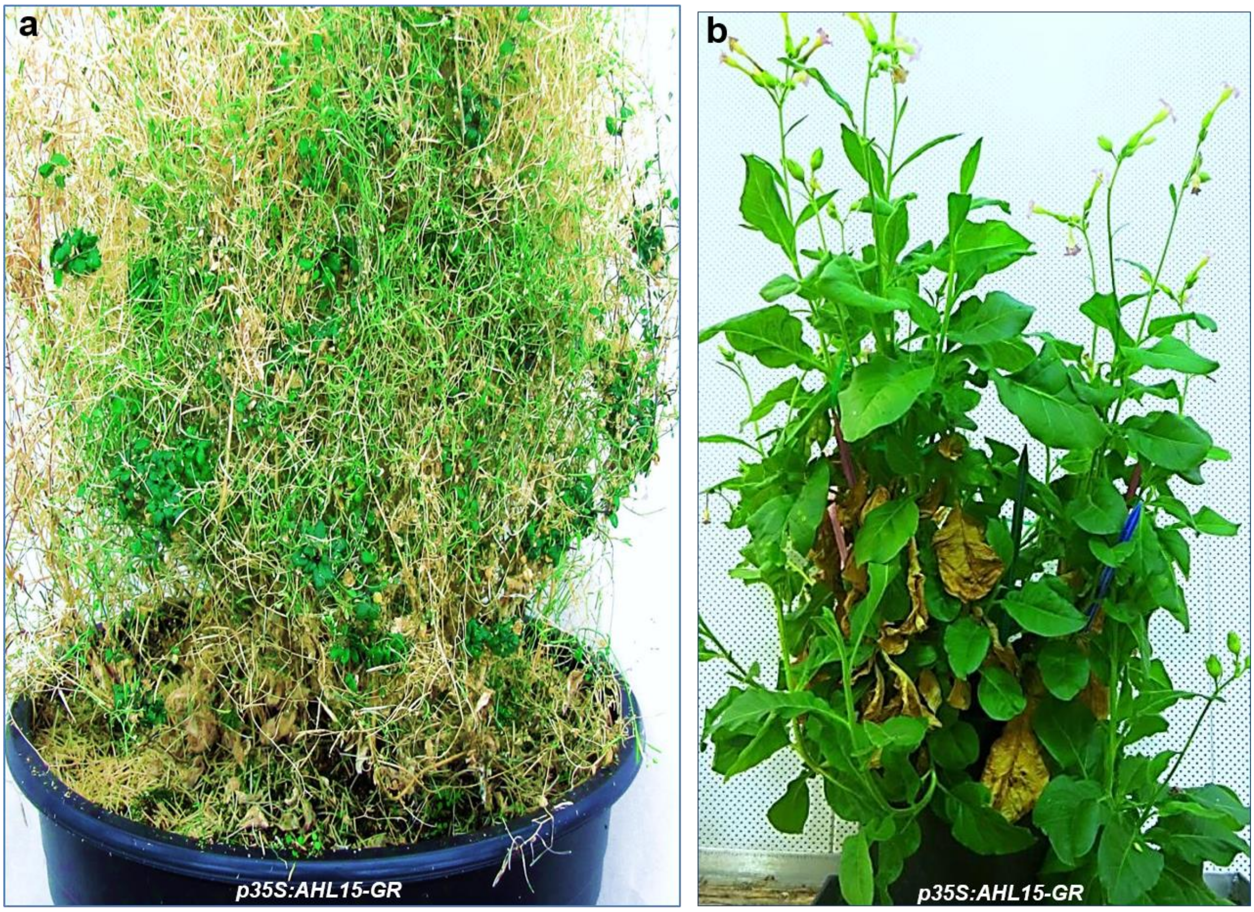


**Supplementary Figure 11 | *AHL15* overexpression promotes longevity in Arabidopsis and tobacco.** (**a**) Renewed vegetative growth on aerial branches of a 5-month-old *35S:AHL15-GR* plant, 4 weeks after spraying with 20 μM DEX. For presentation purposes, the original background of the image was replaced by a homogeneous white background. (**b**) Efficient production of leaves and inflorescences in a 3-year-old *35S::AHL15-GR* tobacco plant, 3 weeks after treatment with 30 μM DEX, following 6 previous cycles of DEX-induced seed production. Plants in **a** and **b** were grown in LD conditions.

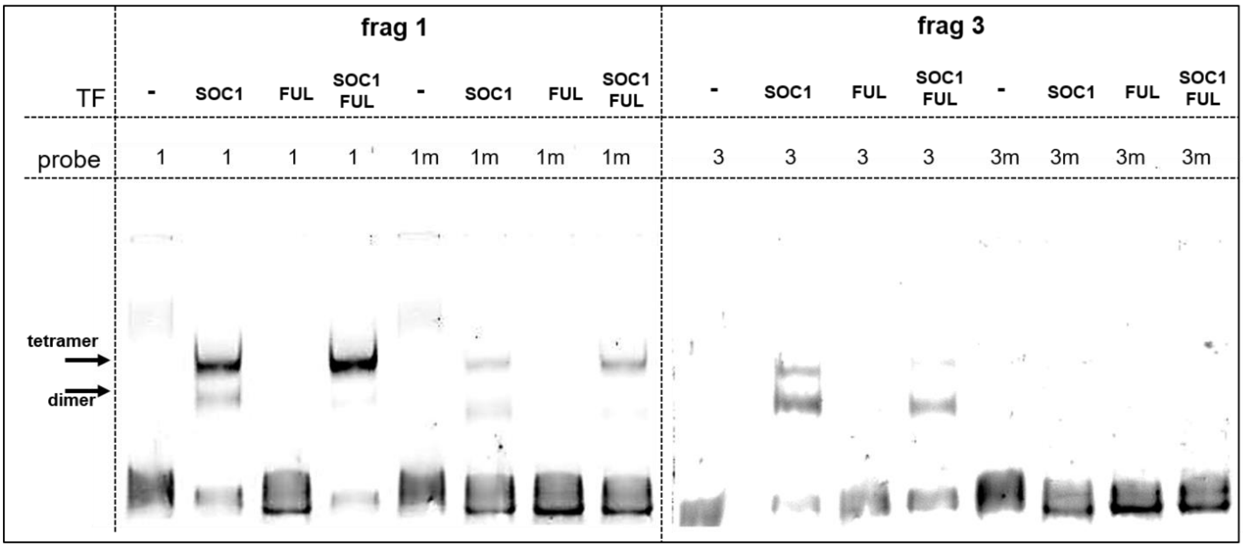


**Supplementary Fig. 12. Binding of SOC1 and FUL to regulatory regions near *AHL15*.** Left panel: EMSA of promoter fragment 1 with a wild-type (1) or mutated (m1) CArG-box. Right panel: EMSA of downstream fragment 3 with a wild-type (3) or mutated (3m) CArG-box. Shifting of the probe, indicating binding, occurs either by a tetramer (upper band) or dimer (lower band).

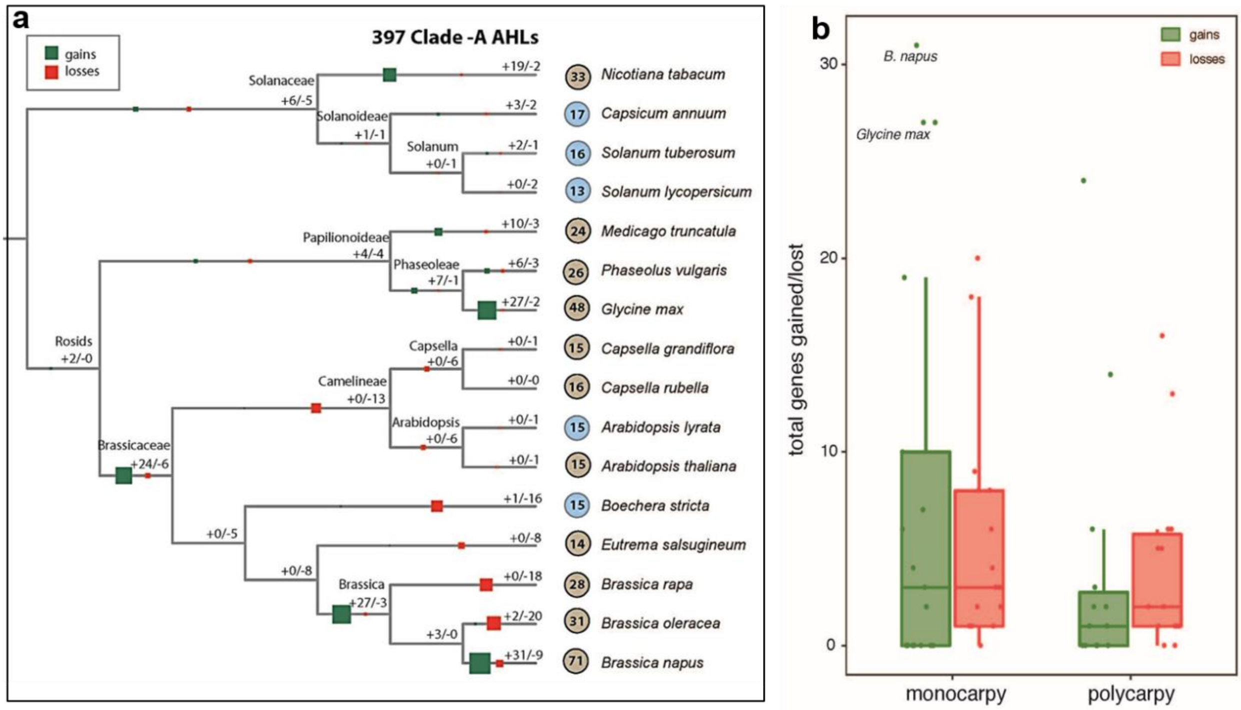


**Supplementary Figure 13. Evolutionary history of the Clade-A *AHL* gene family in relation to the monocarpic and polycarpic plant growth habit.** (**a**) Reconciliation of the gene (protein) tree with the species tree of 16 selected dicot plant species with either a monocarpic (light brown circled number) or a polycarpic (light blue circled number) life history strategy. The tree topology was extracted from the NCBI taxonomy database. For each tree node, the number of gained genes (+) is indicated by the size of the green box, and the number of lost genes (-) by the size of the red box. The total number of *AHL* genes for each plant species is indicated by the circled number. (**b**) The distribution of total Clade-A *AHL* gene gains and losses for monocarpy or polycarpy in (a) have been summarized as box-plots. The character state of each ancestor was determined for the presence (polycarpy) or absence (monocarpy) of polycarpic growth habit using Dollo's parsimony principle. Horizontal bars indicate median values, boxes demarcate 1st and 3rd quartiles, and whiskers delineate minimum (within 1.5 times Inter Quartile Range (IQR) below 1st quartile) and maximum (within 1.5 times IQR above 3rd quartile) values. The dots in the box-plots indicate all the ancestral states and current species for which the total gene gains/losses were estimated. The outlier species are labelled with the species name. The outlier species are labelled with the species name.

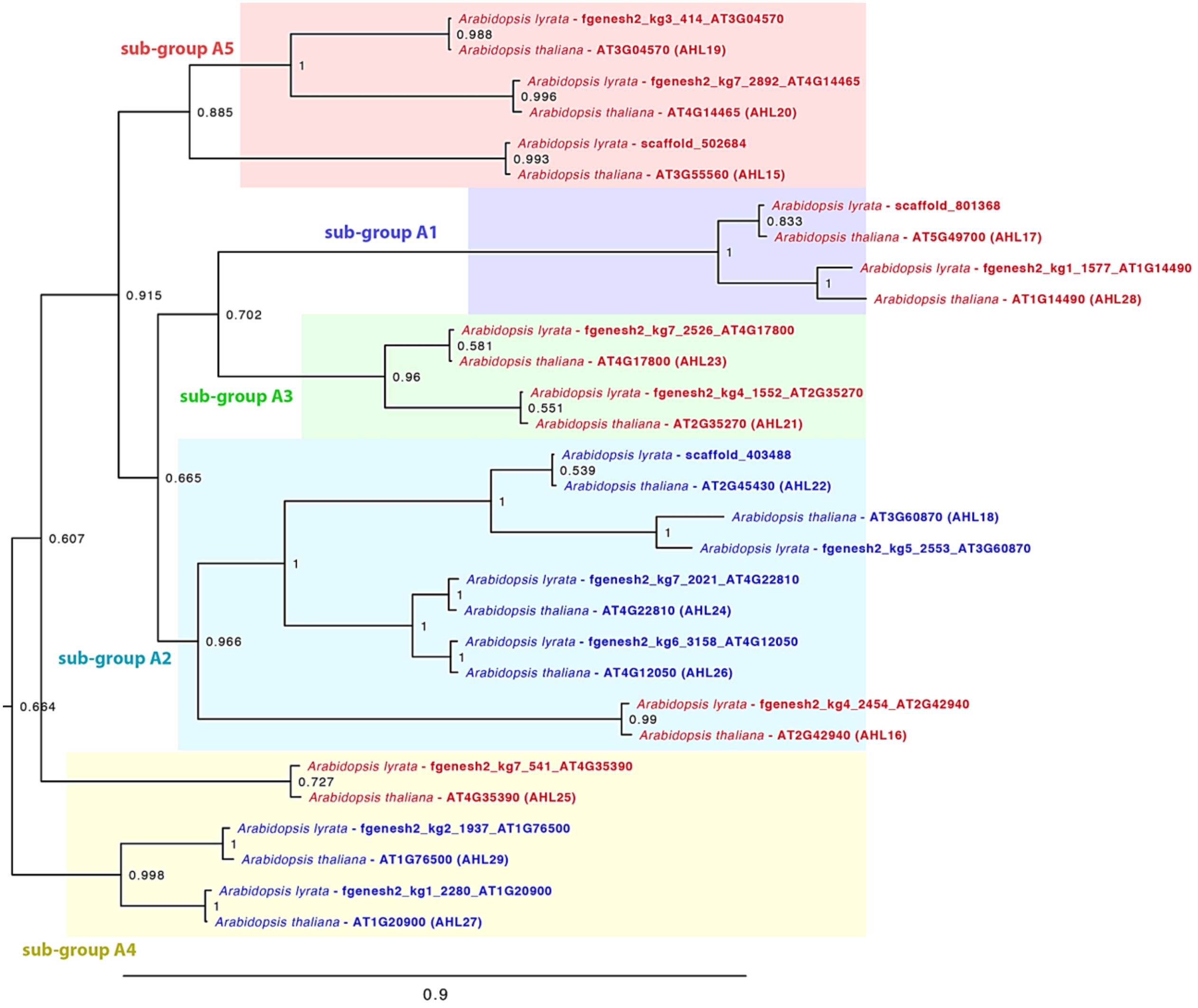


**Supplementary Figure 14 | Phylogenetic tree of *A. thaliana* and *A. lyrata* clade-A AHL proteins.** The tree shows 15 one-to-one orthologous pairs between the two species.

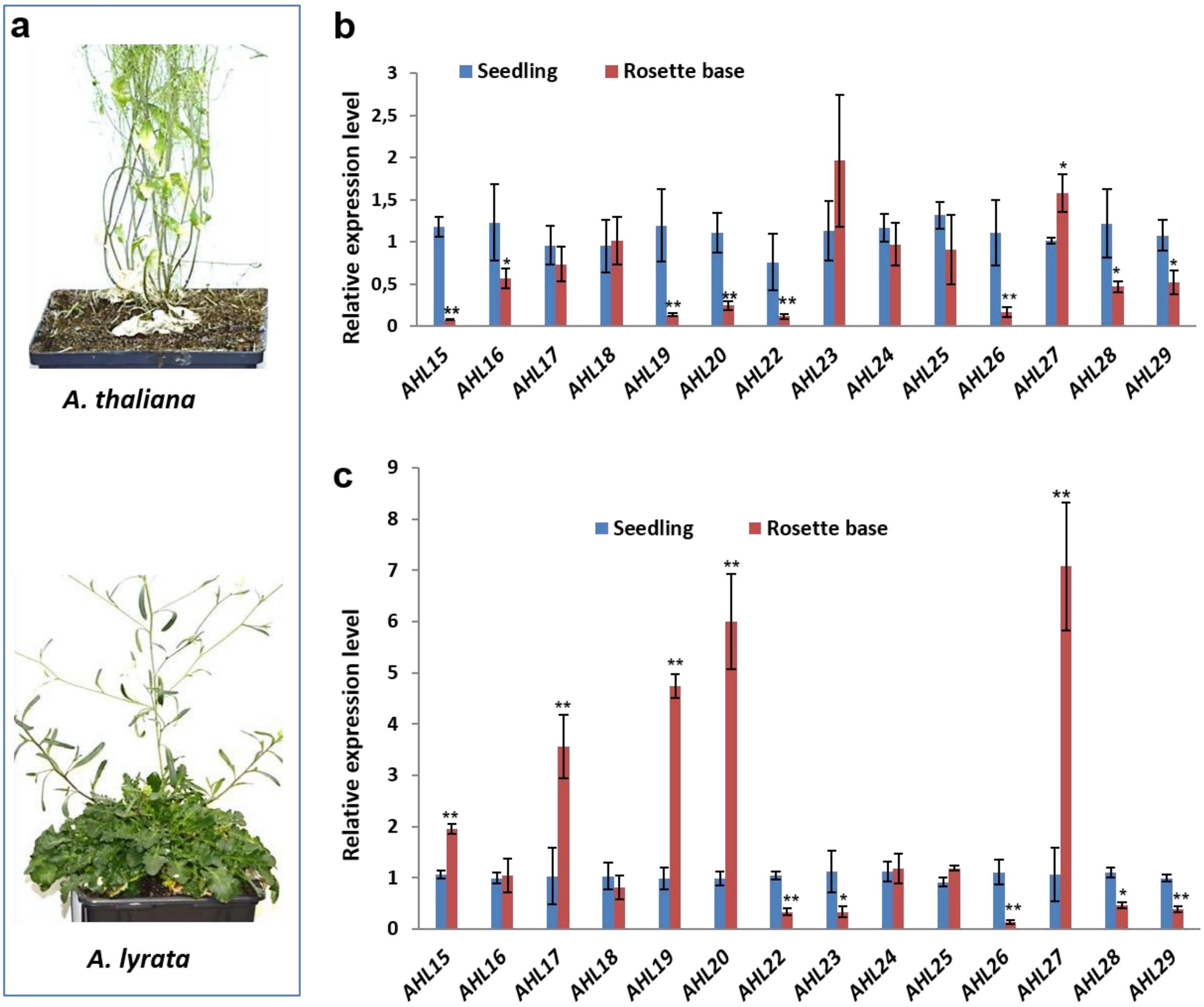


**Supplementary Figure 15 | Expression of clade-A *AHL* genes in seedlings or in the rosette base of flowering *A. thaliana* or *A. lyrata* plants.** (**a**) Shoot phenotype of a 3-month-old *A. thaliana* (upper panel) or a 4-month-old *A. lyrata* (lower panel) plant grown in LD conditions. (**b,c**) qPCR analysis of the expression of clade-A *AHL* genes in 2-week-old seedlings or in the rosette base of 2-month-old flowering plants of *A. thaliana* (**b**) or *A. lyrata* (**c**). Asterisks in **b** and **c** indicate that basal node expression is significantly different from that in seedlings (Student's *t*-test: * p<0.05, ** p<0.01). Error bars represent the standard error of the mean (n =3).

**Supplementary Table 1**: PCR primers used for cloning and genotyping

| Name* | Sequence (5' to 3') | Purpose |
|---|---|---|
| Gateway-AHL15-F | GGGGACAAGTTTGTACAAAAAAGCAGGCTCGATGGCGAATCCTTGGTGGGT | *pGW-AHL15* construct |
| Gateway-AHL15-R | GGGGACCACTTTGTACAAGAAAGCTGGGTAATACGAAGGAGGAGCACGAG | |
| ccdB gene +KpnI-F | CGGGTACCGCTATCGAACCACTTTGTAC | Amplify *ccdB* gene |
| ccdB gene +SphI-R | GACTGCAAGTCACGTCGGCAG | |
| pAHL15:AHL15-F | GGGGACAAGTTTGTACAAAAAAGCAGGCTCGACACTCCTCTGTGCCACATT | *pAHL15:AHL15* construct |
| pAHL15:AHL15-R | GGGGACCACTTTGTACAAGAAAGCTGGGTATCTTTTTTTCTTCTCTAATGG | |
| pFD:AHL15-F | GGGGACAAGTTTGTACAAAAAAGCAGGCTGGCCCTCTCTACTTGATTTAG | *pFD:AHL15* construct |
| pFD:AHL15-R | GGGGACCACTTTGTACAAGAAAGCTGGGTATGGAAAAGAGAACAGAAGTGAAC | |
| pMYB85:AHL15-F | GGGGACAAGTTTGTACAAAAAAGCAGGCTGGTGGGGTGTTGAAATGTCAC | *pMYB85:AHL15* construct |
| pMYB85:AHL15-R | GGGGACCACTTTGTACAAGAAAGCTGGGTATAAATACTATATAGAAATGATATG | |
| pMYB103:AHL15-F | GGGGACAAGTTTGTACAAAAAAGCAGGCTTTCTATTGCTCCTCCTTAAAGG | *pMYB103:AHL15* construct |
| pMYB103:AHL15-R | GGGGACCACTTTGTACAAGAAAGCTGGGTAGATTAGTAGCTCCTCAAAGTAAC | |
| p35S:AHL29-F | ATAAGAATGCGGCCGCGACGGTGGTTACGATCAATC | *p35S:AHL29* construct |
| p35S:AHL29-R | ATAGTTTAGCGGCCGCCTAAAAGGCTGGTCTTGGTG | |
| p35S:AHL20 –F | ATAAGAATGCGGCCGCGCAAACCCTTGGTGGACGAAC | *p35S:AHL20* construct |
| p35S:AHL20-R | ATAGTTTAGCGGCCGCTCAGTAAGGTGGTCTTGCGT | |
| p35S:AHL27-F | ATAAGAATGCGGCCGCGAAGGCGGTTACGAGCAAGG | *p35S:AHL27* construct |
| p35S:AHL27-R | ATAGTTTAGCGGCCGCTTAAAAAGGTGGTCTTGAAG | |
| p35S: BoAHL15-1-F | ATAAGAATGCGGCCGCGCGAATCCTTGGTGGGTAGA | *p35S:BoAHL15* construct |
| p35S: BoAHL15-1-R | ATAGTTTAGCGGCCGCTCAATATGAAGGAGGACCAC | |
| p35S: MtAHL15-F | ATAAGAATGCGGCCGCTCGAATCGATGGTGGAGTGG | *p35S: MtAHL15* construct |
| p35S: MtAHL15-R | ATAGTTTAGCGGCCGCTCAATATGGAGGTGGATGTG | |
| p35S:AHL19-F | GGGGACAAGTTTGTACAAAAAAGCAGGCTCGATGGCGAATCCATGGTGGAC | *p35S:AHL19* construct |
| p35S:AHL19-R | GGGGACCACTTTGTACAAGAAAGCTGGGTAAACAAGTAGCAACTGACTGG | |
| SALK_040729-F | GTCGGAGAGCCATCAACACCA | *ahl15* genotyping |
| SALK_040729-R | CGACGACCCGTAGACCCGGATC | |
| soc1-6 -F | AAAGGATGAGGTTTCAAGCG | *soc1-6* genotyping |
| soc1-6 -R | ATGTGATTCCACAAAAGGCC | |
| ful-7-F | TTTCCGCCTTCTCTCGTTGTG | *ful-7 genotyping* |

*, F: forward; R: reverse

**Supplementary Table 2**: PCR primers used for qRT-PCR and RT-PCR

| Name* | Sequence (5' to 3') | Purpose |
|---|---|---|
| qAHL15-F | AAGAGCAGCCGCTTCAACTA | *qRT-PCR AHL15* |
| qAHL15-R | TGTTGAGCCATTTGATGACC | |
| qAHL16-F | CCAGCTTCATCAGGAGCAAT | *qRT-PCR AHL16* |
| qAHL16-R | ATGCAGCCATGAGAACAACA | |
| qAHL17-F | GTCCACTTATTTCGGCAGGA | *qRT-PCR AHL17* |
| qAHL17-R | TACCCCACACGACTCTCCTC | |
| qAHL18-F | ACACGGACGGTTTGAGATTC | *qRT-PCR AHL18* |
| qAHL18-R | GCCTCTCGTAAGATGCGTTT | |
| qAHL19-F | CTCTAACGCGACTTACGAGAGATT | *qRT-PCR AHL19* |
| qAHL19-R | ATATTATACACCGGAAGTCCTTGGT | |
| qAHL20-F | CAAGGCAGGTTTGAAATCTTATCT | *qRT-PCR AHL20* |
| qAHL20-R | TAGCGTTAGAGAAAGTAGCAGCAA | |
| qAHL22-F | AGCTGGAGCGGTTGCTAATA | *qRT-PCR AHL22* |
| qAHL22-R | CAGCTGGCAATTGAACAGAA | |
| qAHL23-F | TTGTGACGCTACAAGGAACG | *qRT-PCR AHL23* |
| qAHL23-R | AAACGAAGCTGCAATCACAA | |
| qAHL24-F | TGGTTGGAGGAAGCGTAGTT | *qRT-PCR AHL24* |
| qAHL24-R | GCTTGTTGCTGATGTTGCAG | |
| qAHL25-F | GCAAACGCAGTTTATGATAGGTTAC | *qRT-PCR AHL25* |
| qAHL25-R | ATTCCAAGATTGTAGAAAGCAACAC | |
| qAHL26-F | GGTGGGACCTTTGTTGTGTT | *qRT-PCR AHL26* |
| qAHL26-R | TGCCATAGCTTGTTGCTGTC | |
| $_{R1}$AHL15-F | ATGGCGAATCCTTGGTGGGTAG | *RT-PCR$_1$ AHL15* |
| $_{R1}$AHL15-R | TCAATACGAAGGAGGAGCACGAG | |
| $_{R}$ACTIN2-F | TGAGACCTTTAACTCTCCCGCTA | *RT-PCR ACTIN2* |
| $_{R}$ACTIN2-R | TGATTTCTTTGCTCATACGGTCA | |
| $_{R2}$AHL15-F | TCAGCT CCT TCT TTGCACCAC | *RT-PCR$_2$ AHL15* |
| $_{R2}$AHL15-R | ATACGAAGGAGGAGCACGAGG | |
| $_{R3}$AHL15-F | AAGAACAGAACAGCAGAGACG | *RT-PCR$_3$ AHL15* |
| $_{R3}$AHL15-R | TCAATACGAAGGAGGAGCACG | |
| qβ-TUBULIN-6-F | TGGGAACTCTGCTCATATCT | *qRT-PCR AHL15* |
| qβ-TUBULIN-6-R | GAAAGGAATGAG GTTCACTG | |
| QEF1ALPHA-F | TGAGCACGCTCTTCTTGCTTTCA | *qRT-PCR EF1alpha* |
| QEF1ALPHA-R | GGTGGTGGCATCCATCTTGTTACA | |
| ChIP REF1-F | TCTCCGACCTTTCTTCACACCCATTCC | *ChIP-qPCR REF1* |
| ChIP REF1-R | CTGAGAACTTGCTTACTTGATAGACTC | |
| ChIP REF2-F | GCTATCCACAGGTTAGATAAAGGAG | *ChIP-qPCR REF2* |
| ChIP REF2-R | GGACTAGATTTGAGGAAAGGAAGGA | |
| ChIP frag1-F | TGTCACACCACTCTCTTTGCA | *ChIP-qPCR frag1* |
| ChIP frag1-R | AATAATGGTTACTGAAAACGTACA | |
| ChIP exon-F | TCCAGAGCCATGTTCTTGAG | *ChIP-qPCR exon* |
| ChIP exon-R | GCTGACGCAGAGTAACATTAG | |
| ChIP frag3-F | TTGGATCTTTGGCATTGTCTC | *EMSA-PCR frag3* |
| ChIP frag3-F | TGCTACGGAGTTTAGTCATCA | |
| EMSA frag1-F | TGTCACACCACTCTCTTTGCA | *PCR frag1* |
| EMSA frag1-R | AATAATGGTTACTGAAAACGTACA | |
| EMSA exon-F | TCCAGAGCCATGTTCTTGAG | *PCR exon* |
| EMSA exon-R | GCTGACGCAGAGTAACATTAG | |
| EMSA frag3-F | TTGGATCTTTGGCATTGTCTC | *PCR frag3* |
| EMSA frag3-F | TGCTACGGAGTTTAGTCATCA | |
| EMSA frag1 sequence | TGTCACACCACTCTCTTTGCATATAATCCAAAACATATATGGGTTAATGAATTTTC<br>AAAAAATAACAGCCCTTATTTGGATCTTACAATAAATGTACGTTTTCAGTAACCA<br>TTATT | |
| EMSA mutated frag1 sequence | TGTCACACCACTCTCTTTGCATATAATCCAAAACATATATGGGTTAATGAATTTTC<br>AAAAAATAACAGCCCTTCCTTGGATCTTACAATAAATGTACGTTTTCAGTAACCA<br>TTATT | |
| EMSA exon sequence | TTGGATCTTTGGCATTGTCTCTGGTCTTAAATATCCCTTTTATGGTATATCAAAAA<br>AACATGAGTACGGAATATATGGTCCCATGATGACTAAACTCCGTAGCA | |

| | |
|---|---|
| EMSA frag3 sequence | TTGGATCTTTGGCATTGTCTCTGGTCTTAAATATCCCCTCCATGGTATATCAAAAA<br>AACATGAGTACGGAATATATGGTCCCATGATGACTAAACTCCGTAGCA |
| EMSA mutated frag3 sequence | TCCAGAGCCATGTTCTTGAGATTGCTACGGGAGCTGACGTGGCGGAAAGCTTAA<br>ACGCCTTTGCTCGTAGACGCGGCCGGGGCGTTTCGGTGCTGAGCGGTAGTGGTTT<br>GGTTACTAATGTTACTCTGCGTCAGC |

*, F: forward; R: reverse.
